# Characterization of metabolic changes associated with floral transition in Arabidopsis: *RAFFINOSE SYNTHASE 5* contributes to determine the timing of floral transition

**DOI:** 10.1101/2022.04.29.490013

**Authors:** Jesús Praena Tamayo, Ilara Gabriela Frasson Budzinski, Nicolas Delhomme, Thomas Moritz, Francisco Madueño, Reyes Benlloch

## Abstract

Integration of environmental and endogenous cues triggers floral induction at the optimal time during the plant life cycle. Flowering is a tightly regulated process, which involves an intricated genetic network, as expected for a process crucial for plant fitness and survival. Individual metabolites are known to contribute to the determination of flowering time, including carbohydrates and hormones. However, a global analysis of metabolic changes associated with flowering was still lacking. We performed a metabolomic study to characterize global metabolic changes associated with photoperiodic floral induction. By using an inducible system, with the *CONSTANS* (*CO*) promoter driving the expression of CO fused to the rat glucocorticoid receptor (CO::GR), we induce flowering and identify metabolites that increase or decrease in leaves and apices during floral induction. Combining metabolomic with transcriptomic data, we identify that raffinose metabolism was altered in apices that are induced to flower. Loss-of-function mutants affecting *RAFFINOSE SYNTHASE 5* (*RS5*), a key enzyme of the raffinose metabolism, show an early flowering phenotype. Also, *RS5* expression changes during floral transition, suggesting a role for raffinose catabolism on the release of simple sugars at the apex. We propose that variation on the differential accumulation of raffinose and mono- and disaccharides during floral transition contributes to the induction of floral transition, by influencing expression of *THEHALOSE-6-PHOSPHATE SYNTHASE 1* (*TPS1*) and S*QUAMOSA PROMOTER BINDING PROTEIN-LIKE 3* (*SPL3*), which affect expression of the florigen FLOWERING LOCUS T (FT).

## Introduction

Genetic control of flowering time in Arabidopsis has been intensively studied during the last two decades. There is abundant information on the genetic network contributing to perception, transduction and integration of environmental and endogenous cues controlling flowering time (photoperiod, temperature, carbohydrate status, hormones, age and the autonomous pathway) (Andrés & Coupland, 2012; Kinoshita & Richter, 2020). Integration of the photoperiod signal in leaves and the molecular mechanism by which the florigen triggers floral transition in the apex has been extensively studied and recently reviewed in several works (Luo et al., 2021; Shim et al., 2017; You et al., 2017; Zhu et al., 2021). The CONSTANS (CO)-FLOWERING LOCUS T (FT) module is at the center of the molecular mechanisms perceiving the photoperiodic signal. *CONSTANS* codes for a B-box transcription factor, whose expression and stability are strictly regulated during the day, integrating light and circadian clock cues. In Arabidopsis, when *CO* expression coincides with the light period, under long days (LD), CO protein is stabilized and activates expression of the florigen, *FLOWERING LOCUS T* (FT), in specific phloem companion cells. FT travels though the phloem and reaches the shoot apical meristem, where it triggers floral transition by promoting the expression of floral integrators, such as *LEAFY (LFY)* and *SUPRESSOR OF OVEREXPRESSION OF CONSTANS 1 (SOC1)* (Freytes et al., 2021). Detailed characterization of these molecular events has revealed that the florigen and related flowering time genes form an intricated gene regulatory network that modulates many different aspects of plant development (Osnato et al., 2022; Pin & Nilsson, 2012). It is also increasingly clear that the basic functioning of those gene regulatory circuits controlling photoperiodic flowering is conserved even in phylogenetic distant species, although specific regulatory modules are present in different lineages and have been modulated during domestication (Brambilla et al., 2017; Gaudinier & Blackman, 2020; Khosa et al., 2021; Lin et al., 2020).

Comparatively, little information is available on the metabolic changes that occur in leaves and in the shoot apex during floral transition. Recently, characterization of the metabolome in response to nitrogen deficit has identified developmental stage-specific metabolite signatures (Olas et al., 2021). Besides, changes in biosynthesis and degradation of several hormones have been also found to influence flowering in Arabidopsis (Conti, 2017; Izawa, 2020). Several studies have pointed out the relevance of carbohydrate availability for the determination of the timing of flowering. Indeed, the photoperiodic signal has been shown to control carbon distribution during floral transition. On one hand, CO regulates expression of the *GRANULE BOUND STARCH SYNTHASE* (*GBSS*) gene, contributing to sugar mobilization during floral transition (Ortiz-Marchena et al., 2014). On the other hand, FT promotes the expression of *SWEET10* which codes for a sugar transporter active both in vascular tissue of the leaves and in the apex (Andrés et al., 2020). Trehalose-6-phosphate (T6P) has emerged as an important signaling molecule acting as a carbohydrate status sensor, modulating crucial developmental decisions including vegetative and reproductive phase change (Fichtner & Lunn, 2021; Ponnu et al., 2011, 2020; Wahl et al., 2013).

In this context, a metabolomic approach to understand the complex changes that might underly the change of fate in the shoot apical meristem during floral transition is of great interest. The analytical power of metabolomic approaches has greatly improved in the last decade (Patti et al., 2012). Moreover, the application of multi-omic approaches, including metabolomics together with transcriptomics and/or proteomics, or even with high-throughput phenotyping, has shown to be useful for the elucidation of relevant pathways (Clark et al., 2021; Grigoreva et al., 2021; Z. Li et al., 2022; Peng et al., 2022). Here, we present a detailed characterization of metabolic changes associated with photoperiodic floral induction by means of a targeted metabolomic approach combined with transcriptome profiling. The combination of these two “omic” approaches has allowed us to identify pathways that are altered during the process of floral induction. Among others, we found that the raffinose metabolism is perturbed in the apex of plants that have been induced to flower compared to those that remain vegetative.

Raffinose is a trisaccharide composed of galactose, glucose, and fructose and belongs, together with stachyose, verbascose and ajugose, to the raffinose family of oligosaccharides (RFOs). The first step of RFO biosynthesis is the synthesis of galactinol from myo-inositol and UDP-galactose by galactinol synthases. Then, raffinose synthases (RS) transfer a galactosyl moiety from galactinol to sucrose to generate raffinose. Further enzymatic additions of galactosyl moiety will result in higher order RFOs (stachyose, verbascose and ajugose). RFOs have a role as storage and transport carbohydrates but in the last years a much wider role has been described for these sugar molecules. They form part of the plant response to abiotic stress, they accumulate in the seed during maturation and protect the embryo during desiccation, they have been related to seed vigor and bud dormancy in several trees, and intriguingly, it has also been suggested that they could have a role in signal transduction (Sengupta et al., 2015).

In this work, we have identified metabolites that accumulate differentially during floral induction. Our data points out several pathways that are altered and might contribute to the control of flowering time, including several hormone biosynthesis and/or conjugation, amino acid and carbohydrate metabolism. Here, we have focused on the characterization of changes in raffinose metabolism during floral transition. We have characterized gene expression changes in key enzymes involved in raffinose synthesis during floral transition and show that mutations affecting the *RAFFINOSE SYNTHASE 5* gene display early flowering phenotype. Our results suggest that a change in the ratio of raffinose to mono and disaccharides in the shoot apical meristem contributes to floral transition by increasing sugar availability, necessary for reproductive (sink) organ formation. This characterization offers new insights in the biochemical changes undergoing at the shoot apical meristem during floral transition. This study opens new lines of research on how metabolic changes contribute and modulate the genetic network controlling the transition to the reproductive phase in Arabidopsis.

## Materials and methods

### Plant material and growth conditions

Arabidopsis plants were grown in a 2:1:1 mixture of black peat:perlite:vermiculite in 4.1×4.5×5 cm pots. After sowing, seeds were stratified in the dark at 4°C for 3-4 days and the plants were grown in greenhouse cabins with natural light supplemented with cold white light (4600lm) to achieve a photoperiod of 16h, and the temperature was kept constant between 21ᵒC and 23ᵒC. Alternatively, plants were grown in culture chambers with cool white fluorescent light (4600lm) under a long-day photoperiod of 16h light and 8h dark at 21ᵒC or a short-day photoperiod of 8h light and 16h dark. Irrigation was carried out twice weekly with Hoagland No. 1 solution enriched with trace elements. *Arabidopsis thaliana* Col-0 was used as wild type. All *Arabidopsis* mutants used in this work have been previously described or have been retrieved from NASC: *co-10* (SAIL_24_H04) (Laubinger et al., 2006), *ft-10* (GK-290E08)(Yoo et al., 2005), *gs1-1* (SALK_121059C) and *gs2-1* (SALK_101144C) (Jing et al., 2018), *gs3-1* (SALK_201770C; N689428), *gs4-1* (SALK_029719C, N661987), *rs2-2* (GK-024G04; N402284), *rs4-1* (SALK_045237C) (Gangl et al., 2015), *rs5-2* (SALK_085989C)(Egert et al., 2013), *rs5-3* (GK-106F01) (Zuther et al., 2004) and *rs6-1* (SALK_035336C, N653138). Double mutants *gs1-1 gs2-1, rs4-1 rs5-2* and *rs5-2 rs6-1* have been generated in this work. Primers used for genotyping of the mutants can be found in Supplementary Table 7.

### Constructs

Construct containing the *FT* promoter (*pFT*) (6kb) and *CO* promoter (*pCO*) (3kb) and the coding sequence for the corresponding fusion protein (FT-GR or CO-GR) were introduced into the pUC57 plasmid with attL1 and attL2 recombination sites for GATEWAY. Then, inserts were cloned into pGWB601 by LR recombination. Constructs were introduced into Arabidopsis Col-0, *co-10* or ft-10 by the floral dip method(Clough & Bent, 1998).

### Plant phenotyping

Flowering time was measure by scoring total leaf number or by scoring days until bolting (bolting refers to days until the inflorescence reached 1 cm).

Total number of fruits produced by the main inflorescence was scored when the shoot apical meristem arrested its growth. For this purpose, newly produced fruits were scored weekly. To score the number of seeds per fruit, mature fruits were collected between the floral nodes 6-15 and performed the counts using an Olympus SZ60 magnifying glass.

Statistical analysis of phenotyping experiments (flowering time, number of fruits and seeds) and expression level (RT -qPCR data) were performed using a t-test or Analysis of Variance (ANOVA) with post-hoc Tukey Honestly Significance Difference (HSD) test (https://astatsa.com/OneWay_Anova_with_TukeyHSD/).

### Sampling plant material for metabolomic and transcriptomic approaches

We stratified seeds for 3-4 days and transfer them to long-day conditions (16h light/8h dark) for 14 days. Then, plants were treated with dexamethasone solution (30 μM dexamethasone in 0.01% Tween-20) or mock solution (0.01% Tween-20) by spraying the rosette (3 sprays per plant corresponding approximately to ≈260 µl). At this time, the plants had 4-5 leaves. The treatment was done at ZT13 (three hours before lights off). After the treatment, trays were covered for 12 hours. The cover was completely removed at ZT10 next day. All apex and leaf samples were collected between ZT14-16. Leaf samples were collected by cutting the petiole of the youngest fully expanded leaf with fine tweezers and immediately freezing it in liquid nitrogen. The apex of the corresponding plant was then harvested, removing all visible petioles, small leaves, root, hypocotyl, and stem. The apex was immediately frozen in liquid nitrogen. All samples were stored at -80 °C until processing for metabolites extractions. For each treatment (Dexa and Mock) and time point (days), 10-11 mg fresh weight of frozen sample from the apices and 16-17 mg from the leaves were used. For extraction of metabolites for targeted metabolomics (GC-TOF-MS and LC-QTOF-MS) and untargeted metabolomics (LC-QTOF-MS). Metabolites were extracted according to the procedure described in (Gullberg et al., 2004), using 1 ml of the extraction mixture (1:3:1 chloroform: methanol:water v/v), from which we obtained 900 μl of supernatant. Subsequently, 166 μl of the apices and 100 μl of the leaf extractions were dried in SpeedVac and prepared for metabolomics analysis using GC-TOF-MS and LC-QTOF-MS. Quality control (QC) samples were prepared by pipetting 50 μl of each sample, each time and each condition up to a stock of 400 μl.

### Targeted Metabolomic by GC-MS and LC-MS

Samples were derivatized overnight at room temperature by adding 30 μL methoxyamine (15 ng/μL pyridine), followed by 30 μL MSTFA with 1% TMCS for 1 hour, and finally 30 μL methyl stearate (15 ng/μL in heptane) was added before injection into the instrument. The internal standards (IS) were added as described in (Gullberg et al., 2004). The GC-TOF-MS instrument used was an Agilent 7890B gas chromatograph equipped with a 10 m × 0.18 mm fused silica capillary column with a 0.18 μm Rxi-5 Sil MS stationary phase (Restek Corporation, U.S.) and connected to a Pegasus BT time-of-flight mass spectrometer, GC-TOF-MS (Leco Corp., St Joseph, MI, USA). Splitless injections were performed using an L-PAL3 autosampler (CTC Analytics AG, Switzerland). The temperature of the injector was 270 °C, the purge flow rate was 20 ml min^-1^ and the column temperature was held at 70°C for 2 minutes, then raised by 40 °C min^-1^ until 320 °C and remained there for 2 minutes. The transfer line temperature was 250 °C and the ion source was 200 °C. Ions were generated with a 70 eV electron beam at an ionization current of 2.0 mA, and 30 spectra/s were recorded in the mass range m/z 50-800. The accelerating voltage was turned on after a solvent delay of 150 s. All generated spectra files were converted to NetCDF file format and peak alignment, integration and feature identification files were processed using in-house scripts for MATLAB ver. 8.1. To determine the retention time index of the detected compounds, an alkane mixture (C12 - C40) was analysed (Schauer et al., 2005). Metabolite identification was performed manually by comparing mass spectra and retention indexes using NIST MS Search v. 2.0 with in house libraries and NIST98 spectral database.

The LC-QTOF-MS instruments used was an Agilent 1290 Infinity UHPLC system (Agilent Technologies, Waldbronn, Germany) for chromatographic separation with an Acquity UPLC HSS T3, 2.1 × 50 mm, 1.8 µm C18 column in combination with a 2.1 mm × 5 mm, 1.8 µm VanGuard precolumn (Waters Corporation, Milford, MA, USA). 2 μl of the resuspended samples (methanol: water 50%) containing internal standards (Gullberg et al., 2004). Mobile phases A (water containing 0.1% formic acid) and B (75/25 acetonitrile: 2-propanol, 0.1% formic acid) were used as elution buffers for the gradient. For each injection, the initial flow rate was 0-5 ml min^-1^ and the compounds were eluted with a linear gradient of 0.1-10% of B for 2 minutes. The gradient was increased to 99% over 5 minutes and held at 99% for 2 minutes. Then B was gradually reduced until the next injection at a flow rate of 0-5 ml min^-1^. The mass spectrometer was an Aligent 6550 Q-TOF with jet stream electrospray ionization in positive or negative ion mode. The same settings were used for both modes, with capillary voltage of 4000 V in positive or negative mode (Abreu et al., 2020).

### Statistical analysis of metabolomic data

For statistical analysis of all experiments related to the metabolome and lipidome of apices and leaves, we used the MetaboAnalyst 5.0 online platform with the Statistical Analysis module (Pang et al., 2021). Normalization was by pooled sample from the group (group PQN) set to day 0, data transformation was “log transformation (base 10)”, and data scaling was “auto-scaling” (mean-centered and divided by standard deviation of each variable). The sPLS- DA for the analysis of the number of components was performed using the validation method “5-fold CV”. The heat map was made considering all time points with the normalized data as the data source and the standardization of the “autoscale features” using the Ward cluster method. The main representative metabolites were identified by analyzing ANOVA.

### Transcriptome analysis

RNA extraction and DNAse treatment for transcriptome analysis were performed using the RNeasy Plant Mini Kit and RNase-Free DNase Set from Qiagen. Extracted RNA was quantified using NanoDrop ND1000 and its quality was determined in a Bioanalyzer 2100 (Agilent).

Samples were sequenced on NovaSeq6000 (NovaSeq Control Software 1.6.0/RTA v3.4.4) with a 151nt(Read1)-10nt(Index1)-10nt(Index2)-151nt(Read2) setup using ‘NovaSeqXp’ workflow in ‘S4’ mode flowcell. The Bcl to FastQ conversion was performed using bcl2fastq_v2.20.0.422 from the CASAVA software suite. The quality scale used is Sanger / phred33 / Illumina 1.8.

Pre-processing of the data was performed according to the guidelines described here: http://franklin.upsc.se:3000/materials/materials/Guidelines-for-RNA-Seq-data-analysis.pdf. Briefly, the quality of the raw sequence data was assessed using FastQC (http://www.bioinformatics.babraham.ac.uk/projects/fastqc/) v0.11.4. Residual ribosomal RNA (rRNA) contamination was assessed and filtered using SortMeRNA (v2.1; Kopylova et al. 2012; settings --log --paired_in --fastx--sam --num_alignments 1) using the rRNA sequences provided with SortMeRNA (rfam-5s-database-id98.fasta, rfam-5.8s-database-id98.fasta, silva-arc-16s-database-id95.fasta, silva-bac-16s-database-id85.fasta, silva-euk-18s-database-id95.fasta, silva-arc-23s-database-id98.fasta, silva-bac-23s-database-id98.fasta and silva-euk-28s-database-id98.fasta). Data were then filtered to remove adapters and trimmed for quality using Trimmomatic (v0.39)(Bolger et al., 2014) settings TruSeq3-PE-2.fa:2:30:10 SLIDINGWINDOW:5:20 MINLEN:50). After both filtering steps, FastQC was run again to ensure that no technical artefacts were introduced. Read counts were determined using salmon (v0.14.1) (Patro et al., 2017) with non-default parameters --gcBias –seqBias and using the ARAPORT11 cDNA sequences as a reference (retrieved from the TAIR resource) (Berardini et al., 2015). Salmon abundance values were imported into R (v3.6.2; R Core Team 2019) using the Bioconductor (v3.10; Gentleman et al. 2004) tximport package (v.1.12.3) (Soneson et al., 2016). For data quality assessment (QA) and visualisation, read counts were normalized using a variance stabilizing transformation as implemented in DESeq2. The biological relevance of the data - e.g. biological replicates similarity - was assessed using Principal Component Analysis (PCA) and other visualizations (e.g. heatmaps), using custom R scripts, available at https://github.com/nicolasDelhomme/arabidopsis-floral-induction. Statistical analysis of differential expression (DE) of genes and transcripts between conditions was performed in R using the Bioconductor DESeq2 package (v1.26.0) (Love et al., 2014), with the following model: ∼ MGenotype * MDay to account for both the genotype and the day of harvesting. FDR-adjusted p-values were used to assess significance, with a common threshold of 1% used throughout. All expression results were generated using custom scripts in R. Differentially expressed genes (DEGs) determined in the previous step were used for Gene Ontology (GO) enrichment analyses using custom R scripts.

### Pathway enrichment analysis (MetaboAnalyst/Plant Metabolomic Network) for targeted metabolomic data and transcriptomic data

Using the MetaboAnalyst platform (Pang et al., 2021), after statistical analysis, we selected the “Pathway Analysis” option and the same data normalization, transformation, and scaling as described in the “Statistical Analysis (T-Test/ ANOVA)” section. The enrichment method chosen was the global test > Relative-betweemess Centrality > Reference Metabolome > Arabidopsis thaliana (KEGG). Using the Plant Metabolics Network platform (Hawkins et al., 2021), performed an analysis of the enrichment of metabolic pathways with those metabolites that showed a significant difference in abundance with a p-value < 0.05 (t-test) for each of the analysis time points (days 1, 3, and 5) and the comparison of conditions (Dexa vs. Mock). For the results from the transcriptome, we selected the DEGs with a fold change > 0.5 and < -0.5 for day 1 and 3.

### Total RNA extraction, cDNA synthesis and quantitative real-time PCR (RT-qPCR)

Arabidopsis RNA extraction for analysis of the expression level of a gene by RT-qPCR was performed using the “E.Z.N.A. Plant RNA kit” with the DNase treatment on the column (Omega) according to the manufacturer’s instructions. RNA was quantified in a Nanodrop ND1000 system (Applied), and its integrity was determined by 1.5% agarose gel electrophoresis. For cDNA synthesis, we started with 3 µg of Arabidopsis RNA previously isolated and treated with DNase. The transcription was performed using the “SuperScript IV” system and “Oligo dT12-18” (Thermo) according to the manufacturer’s instructions.

Real-time quantitative PCR (RT-qPCR) was used for relative quantification of gene expression. To perform RT -qPCR, specific primers for each gene were designed using Primer 3 Plus software. Efficiency of amplification was tested for each pair of primers. RT-qPCR reactions were performed in a final volume of 10µL containing 2µL of cDNA (0.02ng/µL), 2µL of “Premix PyroTaq Eva Green qPCR Mix Plus” (GMC) (5X) and 0.4 µL of each of the primers (5 µM). All reactions were performed in triplicate and in a “7500 Fast PCR System” thermal cycler (Applied). Relative expression of the gene of interest with respect to the constitutive *TIP41* or *IPP2* gene was calculated by the 2^−ΔΔCT^ method (Livak & Schmittgen, 2001). Primers used for each gene can be found in Supplementary Table 7.

### RNA in situ hybridization

Samples were collected in a FAE solution (50% ethanol, 3.7% (v/v) formaldehyde, 5% glacial acetic acid) and placed under vacuum according to the procedure described in (Ferrándiz et al., 2000) with minor modifications. Primers used to amplify the *RS5* probe can be found in Supplementary Table 7.

### Extraction and sugar quantification of sugars by GC-MS

The analysis of sugars in Col-0 and *rs5-2* mutant was performed in the Metabolomics Platform of the Institute of Plant Molecular and Cell Biology (UPV-CSIC) by derivatization followed by gas chromatography-mass spectrometry. All the samples were collected at ZT14-ZT 16 from plants grown in vitro under LD conditions. For extraction, Col-0 and rs5-2 apices and leaves samples (40 mg fresh weight) were homogenized in liquid nitrogen, and 1400 μL 100% methanol with 60 μL internal standard was added to each sample (Ribitol to 0.2 mg/mL in water). The sample was then incubated at 70 °C for 15 minutes and centrifuged at 14,000 rpm for 10 minutes. The supernatant was transferred to a glass vial, and 750 L of CHCl3 and 1500 L of H2O were added. The mixture was shaken for 15 seconds and centrifuged at 14,000 rpm for 15 minutes. 150 μL of the supernatant (aqueous phase, methanol/water) was dried under vacuum for 3 hours. For derivatization, the dried residues were resuspended in 40 µL of 20 mg/mL methoxyamine hydrochloride in pyridine and incubated at 37°C for 90 minutes. Next, 70 µL of MSTFA (N-methyl-N-[trimethylsilyl]trifluoroacetamide) and 6 µL of a standard mixture for adjusting retention times were added to each sample (mixture of fatty acid methyl esters with 8 to 24 carbons at 3.7% [w/v]) and incubated for 30 minutes at 37°C. 100 µL of each sample was transferred to a chromatography vial and gas chromatography was performed. Gas chromatography and mass spectrometry: 2 μL of each sample was injected in splitless and split 1:10 mode into a 6890N gas chromatograph (Agilent Technologies Inc. Santa Clara, CA) coupled to a Pegasus 4D TOF mass spectrometer (LECO, St. Joseph, MI). Gas chromatography was performed using a BPX35 column (30 m × 0.32 mm × 0.25 μm) (SGE Analytical Science Pty Ltd., Australia) with helium as the carrier gas at a constant flow rate of 2 mL/minute. The liner was set at 230°C. The oven program was set at 85°C for 2 minutes and increased to 360°C with a ramp of 8°C per minute. Mass spectra were recorded at 6.25 spectra per second in the range of m/z 35-900 and an ionization energy of 70 eV. Chromatograms and mass spectra were analyzed using CHROMATOF software (LECO, St. Joseph, MI). Compounds of interest were identified by comparison with the spectra of previously traced standards.

## Results

### A system to analyze metabolic changes associated with floral transition

With the aim of generating a system in which we could readily control floral induction in Arabidopsis by inducing *CO* or *FT* expression in the vascular tissue, we generated constructs where the corresponding endogenous promoters drove the expression of a CO or FT protein fusion to the rat glucocorticoid receptor (GR). Alternatively, the promoter of the *SUCROSE-PROTON SYMPORTER 2* (pSUC2) was used to provide a phloem-specific expression of the fusion proteins. To evaluate which combination of promoter and fusion protein provided us with a robust system to induce floral transition, we evaluated the phenotype of pCO::CO::GR, pSUC2::CO::GR, pFT::FT::GR and pSUC2::FT::GR in wild type and in the corresponding mutant backgrounds (*co-10* or *ft-10*) (Supplementary Table 1). Lines carrying the *CO::GR* transgene displayed a very clear flowering response upon dexamethasone (dexa) treatment, independently of the promoter driving the transgene expression (*pCO* or *pSUC2*). *co-10* mutant plants, with either the pCO::CO::GR or pSUC2::CO::GR, treated with dexa showed an early flowering phenotype compared to those treated with mock, which flowered at the same time as the *co-10* mutant. We observed that expression of the CO::GR protein in the wild type background (Col-0) caused an unexpected late flowering phenotype in mock-treated plants as compared to Col-0 plants. We did not explore the molecular basis of this phenotype, since it was not observed in the mutant lines that were used in our studies. We did not observed any response upon dexa treatment in any of the lines expressing the FT::GR protein. We therefore conclude that expressing the CO::GR protein fusion under the endogenous *CO* promoter or the *SUC2* promoter constitutes a robust system to control floral induction. To choose which of the two promoters provided a better response, we evaluated the flowering response after dexa/mock treatment in five T3 single-insert homozygous transgenic lines carrying the pCO::CO::GR or the pSUC::CO::GR trangenes (Supplementary Table 2). We observed that lines carrying the pCO::CO::GR transgene responded better to the dexa treatment, flowering almost as early as Col-0 plants. On the other side, mock-treated plants of the same lines flowered late, as *co-10* plants, confirming that the CO::GR protein in the cytoplasm did not altered photoperiodic flowering in these plants. Among the pCO::CO::GR, line #9 provided a strong response to the dexa treatment, flowering with 25 total leaves compared to the 18 leaves produced by Col-0 and 46 leaves by the *co-10* mutant. Therefore, we selected pCO::CO::GR #9 line for further studies. The *CO* promoter used in this construct specifically drove the expression of the fusion proteins in the vascular tissue, as shown by the expression pattern of the pCO::GUS transgene in the reporter line (Supplementary Figure 1A). Besides, the CO::GR transgene followed the temporal expression pattern described for *CO* under LD conditions, with a low expression during the morning that rises around *Zeitgeber Time* 12 (ZT12) to reach a maximum towards the end of the day (Supplementary Figure 1B). We concluded that the selected line, in addition to efficiently responding to dexa treatment, which induce flowering, replicated both the spatial and the temporal expression of the endogenous *CO* gene, and therefore it is a useful tool to identify key metabolic changes during floral transition.

To determine which was the optimal timing to collect leaf and apex samples for a metabolomic study on the floral induction process, we scored flowering time and upregulation of floral marker genes upon dexa/mock treatment of the pCO::CO::GR #9 line. Plants were grown for 14 days under LD conditions and then treated with dexa or mock. As previously mentioned, pCO::CO::GR plants treated with mock behaved as *co- 10* plants, flowering with more than 40 total leaves. In contrast, dexa-treated plants flowered much earlier, similar to Col-0 (Figure 1A). Thus, upon dexa-treatment these plants initiated floral induction in contrast to the mock-treated ones that remained vegetative for a longer period (Figure 1B). Floral induction was triggered by temporal *FT*-induction in dexa-treated plants (Figure 1C), reaching a maximum expression level one day after treatment. In the apex, upregulation of *SUPRESSOR OF CONSTANS OVEREXPRESSOR 1 (SOC1)* expression confirmed the initiation of the floral transition in dexa-treated plants, further confirmed by upregulation of floral meristem marker genes such as *APETALA 1* (*AP1)* and *LEAFY* (*LFY)* at day 5 after dexa treatment. *TERMINAL FLOWER 1* (*TFL1)* expression was also upregulated at day 3 and the difference with mock-treated plants became evident also at day 5. All the analysed genes displayed a lower expression level at all time points in mock-treated plants, in agreement with those plants being in a vegetative stage (Figure 1D-G). We therefore concluded that, in our experimental conditions, floral transition occurs in dexa-treated plants between day 1 and day 3 and that by day 5 after treatment those plants have undergone floral transition and floral meristems are being produced at the flanks of the meristem.

**Figure 1.**
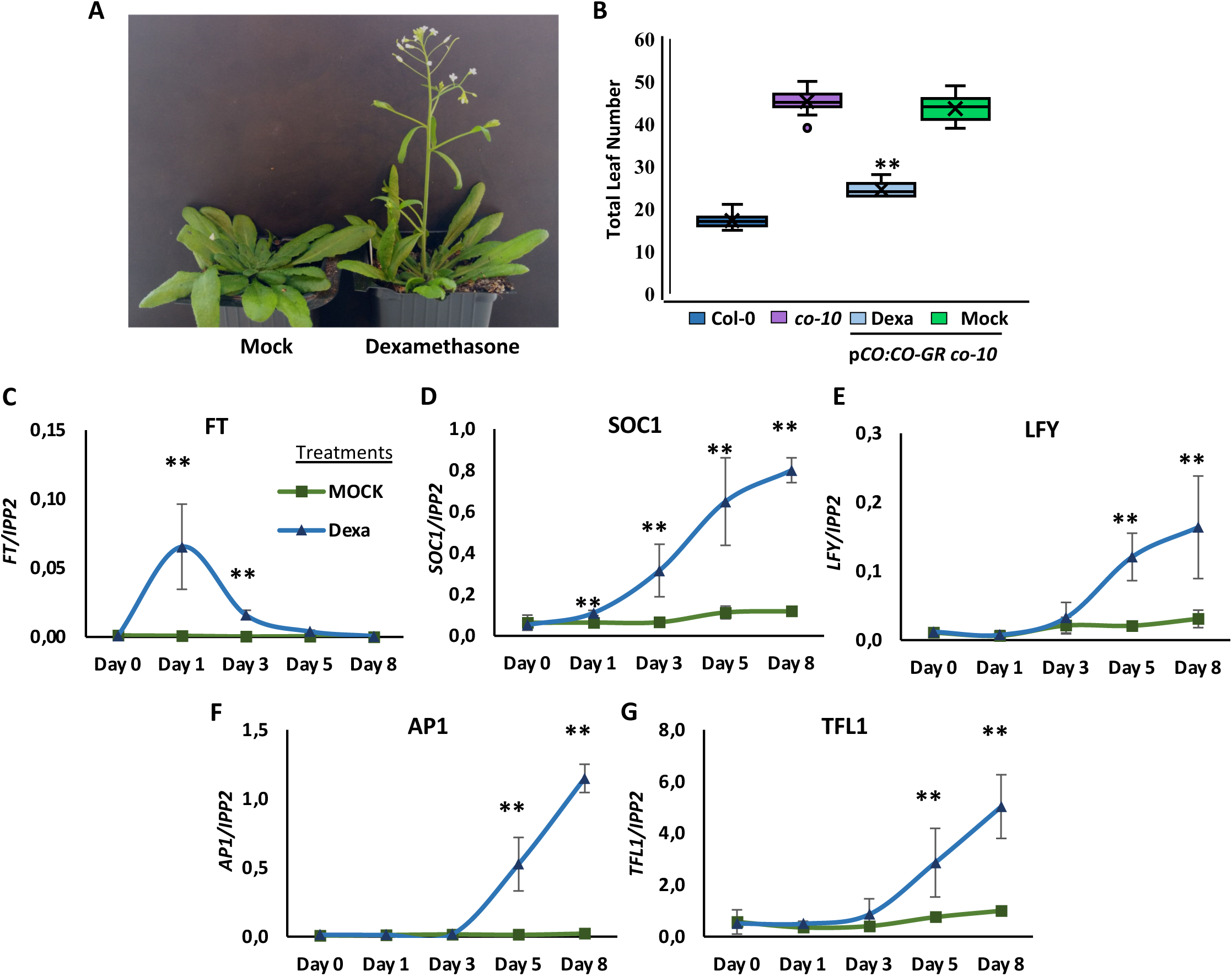
Characterization of flowering time and molecular response to dexamethasone and mock treatment in the selected line p*CO*::*CO-GR co-10* #9. **A.** Phenotype of 42-day old mock- and dexamethasone-treated p*CO*::*CO-GR co-10* #9 plants, grown in LD conditions. B. Flowering time was measured as total leaf number (average ± SD; n=25). **C-G.** Gene expression level of flowering-related genes (*FT*, *SOC1*, *LFY*, *AP1* and *TFL1,* respectively) in response to dexa or mock treatment in the line p*CO*::*CO-GR co-10* #9. Expression levels are the average of three biological replicates, error bars correspond to the SD. IPP2 was sued as a reference gene. ANOVA with Tukey correction was performed to calculate the significant differences. ** indicates p-value<0.01,* indicates p-value<0.05.

### Targeted metabolomics to identify changes in metabolite abundance associated with floral transition

To characterize metabolic changes associated with floral transition, we grew pCO::CO::GR plant for 14 days in LD conditions and then treated them with mock or dexa. Shoot apex and leaf samples were collected at the day of the treatment, and then after 1, 3 or 5 days. Samples were extracted and analysed by Gas Chromatography-Mass Spectrometry (GC-MS) and Liquid Chromatography-Mass Spectrometry (LC-MS). In leaf samples, a total of 65 metabolites were identified by GC-MS and 95 by LC-MS (41 in positive mode and 54 in negative mode). Similarly, 64 metabolites were identified in apex samples by GC-MS and 110 by LC-MS (57 by positive and 53 by negative mode). After removing duplicated metabolites, we identified a total of 136 metabolites in leaf samples and 133 in apex samples (Supplementary Table 3).

Surprisingly, we did not observe any significant differences in metabolite abundance in leaves at different timepoints. However, we did find significant differences in metabolite abundance in apex samples. By applying the Sparse Partial Least Square-Discriminant Analysis (sPLS-DA) algorithm (see Materials and methods section), we identified the three major principal components that explain 76.4 % of the data variance and allow a clear separation between groups (Figure 2A). To achieve a global view of metabolite abundance variation in apex samples, we plotted those metabolites displaying more significant changes across the experiment (Figure 2B). According to the data, there are important metabolic changes between plants induced to flower and those that remain vegetative. We found 25 unique metabolites showing significant differences among dexa and mock treated samples on day 1, 16 metabolites on day 3 and 115 on day 5 (t-test, p-value <0.05) (Supplementary Table 4), when, according to flowering gene markers, transition to flowering has occurred in our conditions (Figure 1).

**Figure 2.**
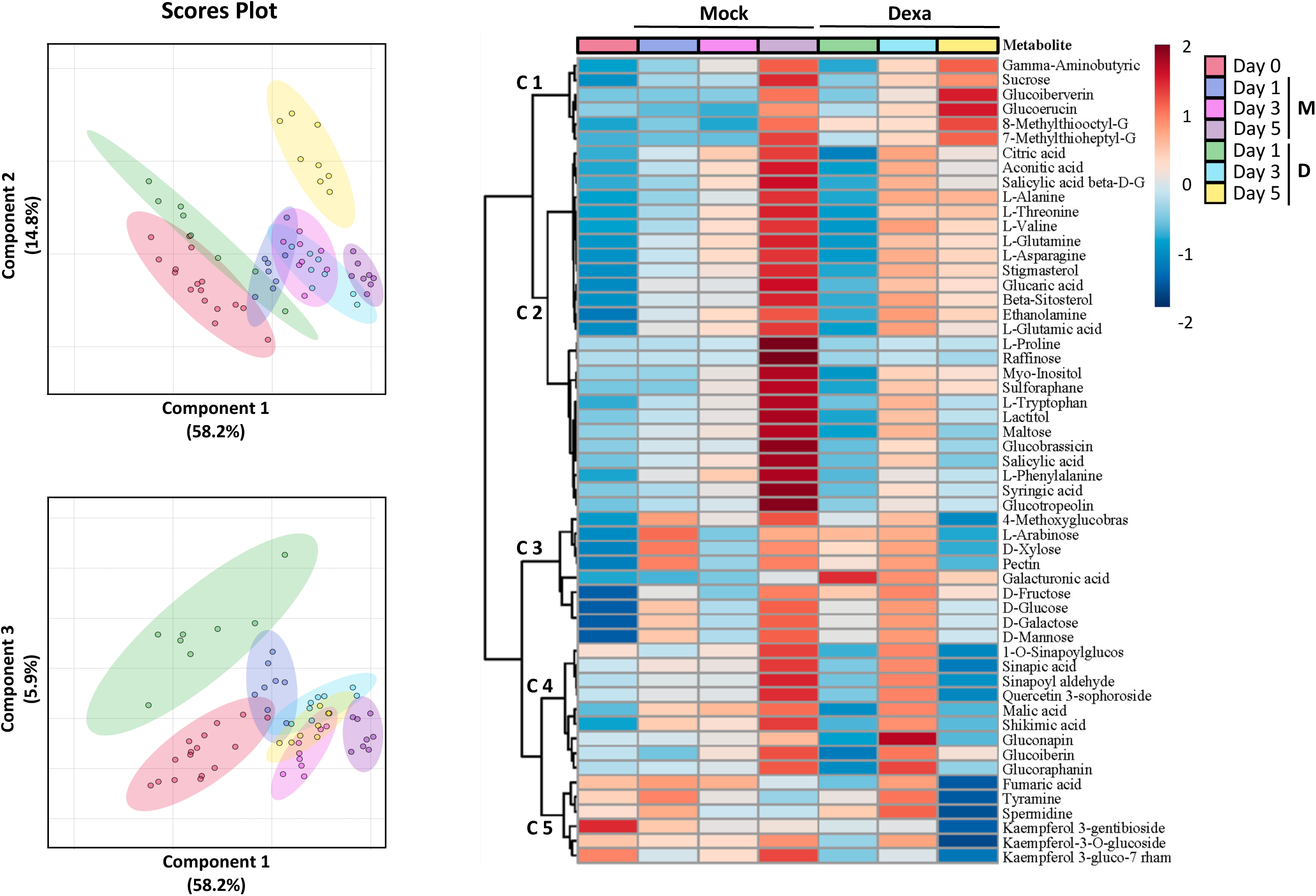
Representation of global metabolic changes in apex samples upon dexa-or mock-treatments. **A.** Apex sample treatments representation through days using the major three principal components (PCs) from sPLS-DA. Score plots between the three major selected PCs. The explained variances are shown in colored brackets assigned per group. **B.** Hierarchical clustering of metabolites represented as a heatmap performed with the top 55 more significant metabolites in all conditions. The ANOVA test with Tukey’s Honestly Significant Difference (Tukey’s HSD) was performed for the multigroup analysis. 0, refers to samples collected before treatments, M refers to the mock-treated samples and D refers to dexamethasone treated samples. The numbers followed by M and D indicate the sampling day after treatment. C1-C5 refers to metabolite clusters. The variables are ranked by the absolute values of their loadings.

We observed a very distinct response in dexa-treated plants between day 1 and day 3, identifying five major clusters of metabolites (Figure 2B; Supplementary Figure 2). Cluster 1 comprises metabolites that increased substantially on mock-treated plants on day five, while abundance in dexa-treated samples remain constant or decreased. Among those we find maltose and raffinose (di and trisaccharide, respectively) (Supplementary Figure 3), salicylic acid (SA), amino acids (phenylalanine, tryptophane and proline) and glucosinolates (glucobrassicin and glucotropeolin), among others. The signalling molecule myo-inositol clustered also in this group, displaying the same tendency in mock and dexa-treated samples but with an attenuated accumulation in the dexa-treated ones. Cluster 2 groups metabolites that remain constant in mock-treated samples until day 5 when their abundance increases and show a continuous tendency to increase in dexa-treated samples, leading to a marked accumulation on day 3 compared to mock. This is the pattern displayed by the monosaccharide sucrose and several glucosinolate metabolites. Metabolites of cluster 3 increase steadily in mock-treated samples while they show a decrease in day 1 and day 5 in dexa-treated samples. Several amino acids and sterol molecules belong to this group, as well as a SA-glucoside. Cluster 4 groups molecules that show an accumulation in dexa samples on day 3 and/or day 5, which is the case of several monosaccharides (glucose, fructose, galactose, mannose, sorbose and arabinose) (Supplementary Figure 3) and an oxidized form of galactose (galacturonic acid). Finally, cluster 5 includes metabolites that decrease their abundance markedly on day 5 in dexa-treated samples, including several flavonoids (kaempferol derivatives), organic acids and glucosinolates. Among all these changes, we identified that there was a different trend regarding the carbohydrate accumulation: on day 1 the trisaccharide raffinose is lower in dexa treated samples while on day 3 we observe an increase in mono- and disaccharides in the same samples. These data suggests that there is a mobilization of storage carbohydrates that impacts on the release of simple sugars at the shoot apex during the process of floral transition (Supplementary Figure 3).

Based on these data, we used two platforms to identify metabolic pathways that change significantly during floral transition: Metaboanalyst (Pang et al., 2021) and Plant Metabolic Network (Hawkins et al., 2021). We focused our analysis on day 1 and day 3, since molecular characterization showed that on day 5 floral transition has already occurred. First, by using the Metaboanalyst platform, we identified 33 pathways that changed significantly among treatments at day 1 (FDR < 0.05) (Supplementary Table 5). Surprisingly, running the same analysis with the dataset corresponding to day 3 we only identified one pathway corresponding to glucosinolate biosynthesis. Secondly, we used all metabolites with significant changes to perform a pathway enrichment study using the Plant Metabolic Network database. In this way, we identified 14 pathways on day 1 and two pathways on day 3 with a p-value <0.01 (Supplementary Table 5). Table 1 summarizes the most relevant results from both analyses. Results revealed that already at day 1 there is a major metabolic adjustment in those apices that have been induced to flower compared to those that remained in a vegetative state. Both analyses indicate alteration in primary metabolic pathways such as the TCA cycle, indicating that the energy requirements at the apex in the vegetative and reproductive stages is different and that mobilization of resources is involved in floral transition. At the same time, we identified an alteration of two carbohydrate pathways, galactose metabolism and stachyose biosynthesis. These two pathways are related since galactose monomers are required for the synthesis of stachyose, a storage oligosaccharide from the raffinose family of oligosaccharides (RFOs). Interestingly, inositol signaling, which is also involved in the synthesis of stachyose, was also altered. Finally, we observed major changes in various amino acid biosynthesis and degradation pathways. In parallel, we also observed perturbation of hormone conjugate synthesis, in particular formation of auxin and jasmonic acid derivatives by conjugation with amino acids. These results suggest that the balance between free and storage carbohydrates and hormone activation/inactivation by conjugation could have a role in early events of floral transition. To further explore this hypothesis, we performed a transcriptomic study in leaf and apex samples with the aim at detecting changes in gene expression of key enzymes acting in altered pathways.

**Table 1.** Analysis of perturbed or enriched pathways using identified metabolites with significant changes in apex samples at day 1.

| **Perturbed pathways - Metaboanalyst** | **FDR** |
| --- | --- |
| Phenylalanine metabolism | 3.71E-03 |
| **Phosphatidylinositol signaling system** | **5.12E-03** |
| Alanine, aspartate and glutamate metabolism | 1.19E-02 |
| Citrate cycle (TCA cycle) | 1.19E-02 |
| Butanoate metabolism | 1.19E-02 |
| Glyoxylate and dicarboxylate metabolism | 1.19E-02 |
| **Galactose metabolism** | **1.41E-02** |
| Arginine biosynthesis | 1.46E-02 |
| Phenylalanine, tyrosine and tryptophan biosynthesis | 1.57E-02 |
| Glycine, serine and threonine metabolism | 2.10E-02 |
| **Inositol phosphate metabolism** | **2.16E-02** |
| Phenylpropanoid biosynthesis | 2.16E-02 |
| Arginine and proline metabolism | 2.27E-02 |
| Glycerophospholipid metabolism | 3.11E-02 |
| Tyrosine metabolism | 3.66E-02 |
| Steroid biosynthesis | 4.39E-02 |
| Glycerolipid metabolism | 4.39E-02 |
| Glutathione metabolism | 5.00E-02 |
| **Enriched pathways - Plant Metabolic Network** | **p-value** |
| Proteinogenic Amino Acid Biosynthesis | 1.53E-06 |
| **Indole-3-acetate conjugate biosynthesis** | **2.13E-06** |
| Amino Acid Biosynthesis | 2.27E-05 |
| **Jasmonoyl-amino acid conjugate biosynthesis** | **2.96E-04** |
| Inactivation | 4.97E-04 |
| Proteinogenic Amino Acid Degradation | 5.23E-04 |
| L-glutamine Biosynthesis | 1.35E-03 |
| Glyoxylate cycle and fatty acid degradation | 1.65E-03 |
| **Stachyose biosynthesis** | **1.75E-03** |
| TCA cycle | 2.19E-03 |
| L-asparagine biosynthesis | 2.78E-03 |
| L-phenylalanine Degradation | 4.04E-03 |
| Glyoxylate cycle | 4.75E-03 |
| Glycine Biosynthesis | 4.75E-03 |

### Transcriptomic changes during floral transition

We conducted a differential gene expression analysis at day 1 and day 3 to characterize transcriptional variations during the floral transition in our experimental system. In leaf samples, we could only identify few genes with an altered expression when comparing dexa *versus* mock treated plants. A total of 24 genes displayed differential expression on day 1 and just one gene on day 3 (log_2_ fold change > 0.5 and FDR < 0.1) (Supplementary Table 6). As expected, *FT* expression was induced in dexamethasone treated leaves, with a massive fold change of 8, confirming that the inducible system worked. Surprisingly, the only gene upregulated in day 3 was *BROTHER OF FT AND TFL1* (*BFT*) with a fold change of 6 (Supplementary Table 6). These few changes at the gene expression level agree with the results obtained for the leaf metabolome, where we did not identify significant differences between treatments.

In apex samples, a total of 571 genes at day 1 and 591 genes at day 3 were differentially expressed comparing samples from dexa and mock treated plants (log_2_ fold change > 0.5 and FDR < 0.05) (Supplementary Table 6). 17 % of all the differential expressed genes (DEGs) encoded enzymes and from those 59% and 43% were downregulated in samples from dexa-treated plants in day 1 and day 3, respectively (Figure 3A).

**Figure 3.**
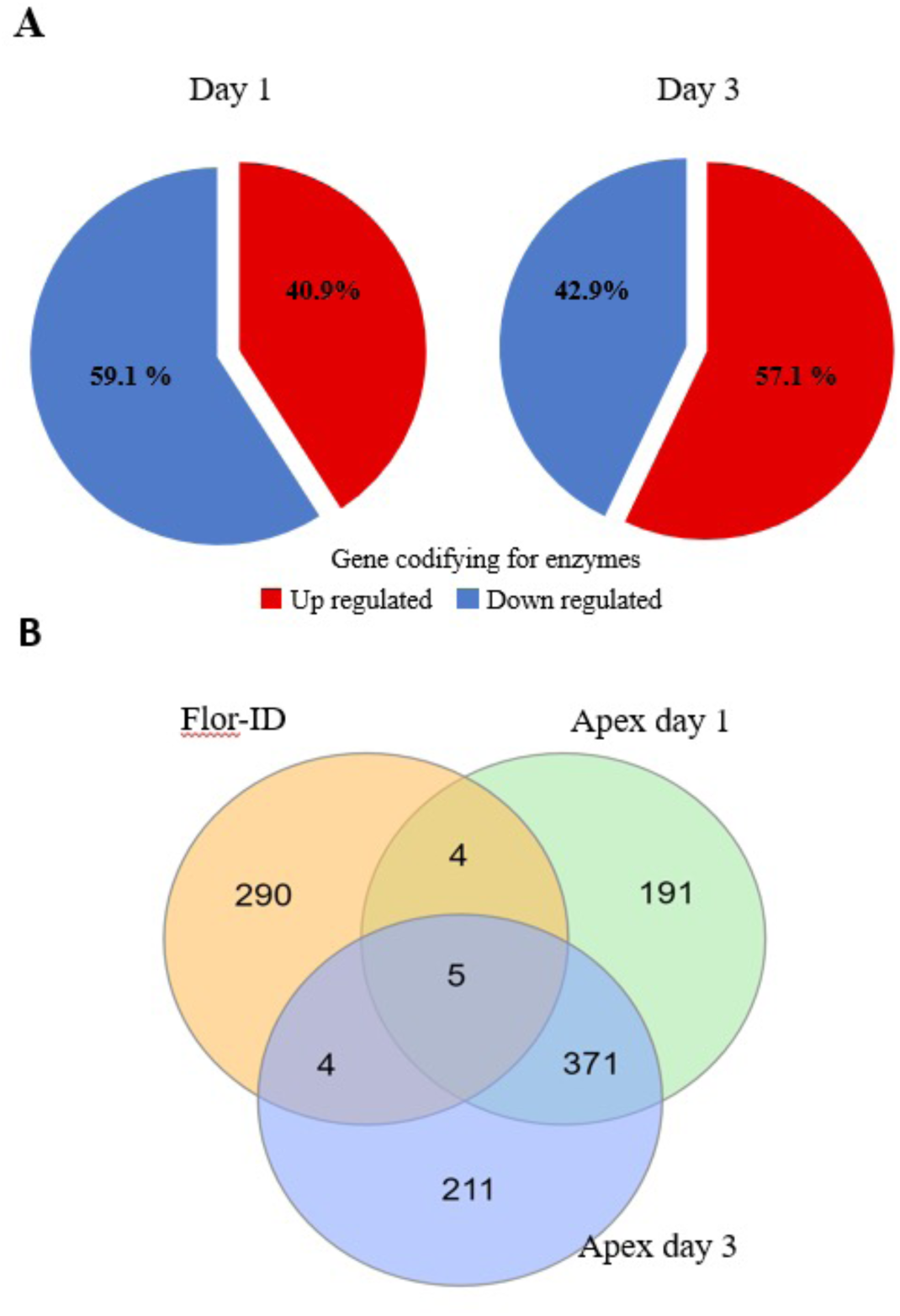
Summary of distribution of genes differentially expressed (DEGs) in apices samples from dexamethasone and mock treated plants. **A.** Percentage of DEGs codifying for enzymes in apex tissue on days 1 and 3. **B.** Venn diagram of RNA-seq data compared with Flor-ID gene database.

To investigate if those transcriptional changes correlate with what has already been described for the floral induction process, we compared the DEG list with the Flowering Interactive Database FLOR-ID (Bouché et al., 2015), which includes gene networks with a known function in the regulation of flowering time (Figure 3B). We detected changes which indicated that dexa-treated apices had initiated floral transition such as the upregulation of *SOC1* and *FRUITFULL* (*FUL*) or the downregulation of the floral repressors *SCHLAFMUTZE* (*SMZ*) and *TARGET OF EAT 3* (*TOE3*). In addition, we could identify other flowering related genes with significant upregulation, including *SQUAMOSA PROMOTER-BINDING-LIKE 4* (*SPL4*), or downregulation, such as *CRYPTOCHROME-INTERACTING BASIC-HELIX-LOOP-HELIX 1* (*CIB1*), *GA INSENSITIVE DWARF 1A* (*GID1A*), *FLOWERING BHLH 4* (*FBH4*), *REVEILLE 2* (*RVE2*), *CONSTANS-LIKE 1* (*COL1*) and *GIBBERELLIN 2-OXIDASE 2* (*GA2OX7*).

We found several genes related to stachyose synthesis strongly downregulated in dexa-treated samples, including *GALACTINOL SYNTHASE 1* (*GOLS1*), *GALACTINOL SYNTHASE 2* (*GOLS2*), *GALACTINOL SYNTHASE 4* (*GOLS4*) and *RAFFINOSE SYNTHASE 2*/*SEED IMBIBITION 2* (*RS2*/*SIP2*). These changes agree with the accumulation of raffinose on mock treated apices as compared to dexa-treated ones, observed in the metabolomics data. On the other hand, myo-inositol is a key compound required for the synthesis of raffinose from galactinol. We identified several genes related to inositol metabolism that were up-or downregulated in dexa-treated apices, including two downregulated genes encoding for phosphatidylinositol phospholipases C (*ATPLC1G* and *PLC4*) and two upregulated genes corresponding to a phosphatidyl-inositol kinase and a phosphatidyl-inositol phosphotransferase (*P4KG4* and *PCS1*). All these transcriptomic data, together with the described changes in metabolites, support the hypothesis that changes in raffinose/stachyose biosynthesis and inositol metabolism are involved in early events of floral transition. Figure 4 summarizes the alterations in raffinose metabolism that we were able to detect by metabolomic or transcriptomic approaches. These results prompted us to focus our efforts to understand the role or contribution of the raffinose pathway to the initiation of floral transition. To do so, we investigated whether loss-of-function mutants affecting stachyose and raffinose biosynthesis displayed a flowering phenotype.

**Figure 4.**
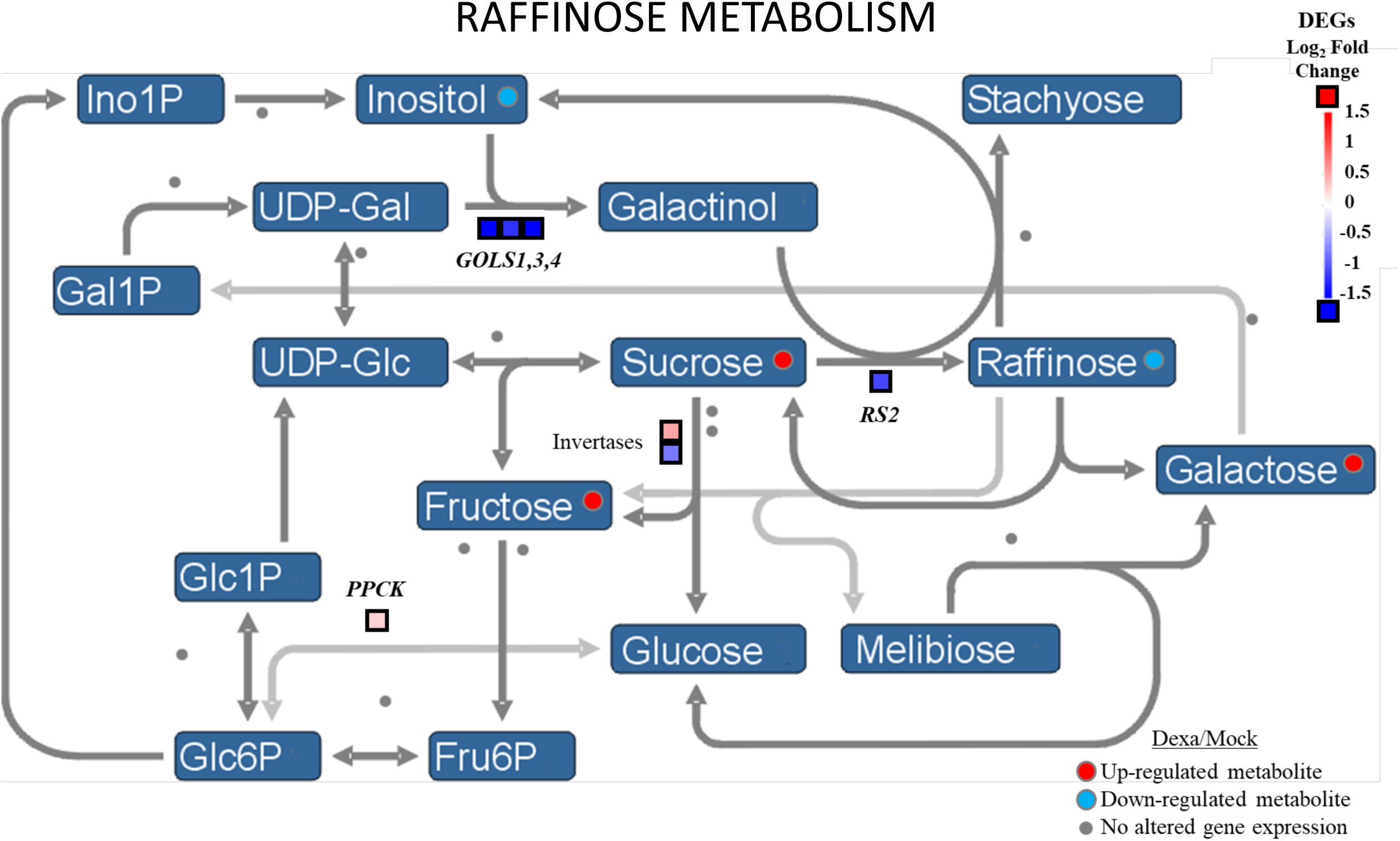
Schematic representation of raffinose metabolism and components of that metabolism detected in the transcriptomic analysis and targeted metabolomic approaches. In the early transition stage (day 1 after dexa treatment), myo-inositol and raffinose decrease and expression of *GOLS1*, *GOLS2*, *GOLS3* and *RS2* is reduced. We also detected changes in the expression of two invertases: *ALKALINE/NEUTRAL INVERTASE C* (AT3G06500; upregulated) and *ALKALINE/NEUTRAL INVERTASE I* (AT4G09510; downregulated). In the late transition stage (day 3 after induction), the expression of *GOLS4*, *GATL10* and *RS2* remain altered. Sucrose, fructose and galactose levels increase. The figure is a modification from the raffinose pathway (modified from MapMan).

### Characterization of loss-of-function mutants affecting raffinose metabolism

We evaluated flowering time in a set of mutants affecting two steps in the raffinose metabolism, galactinol synthesis (*gs1-1*, *gs2-1*, *gs3-1* and *gs4-1*) and raffinose synthesis (*rs2-1*, *rs4-1*, *rs5-2*, *rs5-3* and *rs6-1*). Anticipating the possibility that these enzymes could act redundantly, we also generated and evaluated the phenotype of double mutant combinations (Figure 5A; Table 2). Only mutations affecting *RS5* exhibit an early flowering phenotype. Two independent *rs5* alleles displayed an early flowering phenotype, although in the case of the *rs5-2* allele this phenotype was stronger. Double mutants carrying the *rs5-2* alleles together with *rs2-1*, *rs4-1* or *rs6-1* flowered at the same time as the *rs5-2* mutant.

**Figure 5.**
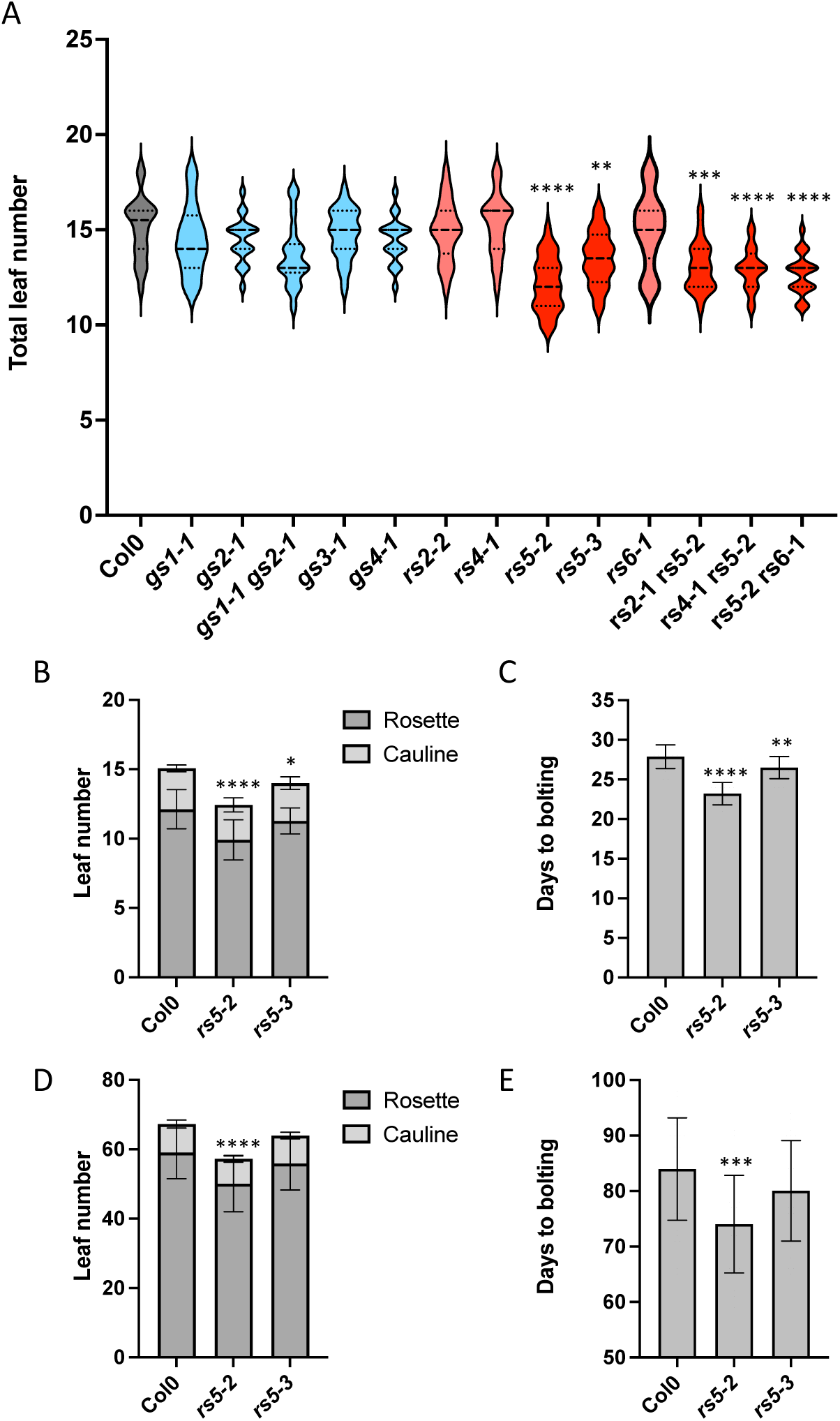
Flowering time characterization of simple and double mutants belonging to the *GOLS* and *RS* gene families. **A.** Flowering time of Col-0, simple mutants *gs1-1, gs2-1, gs3-1, gs4-1, rs2-2, rs4-1, rs5-2, rs5-3, rs6-1* and double mutants *gs1-1 gs2-1, rs2-2 rs5-2, rs4-1 rs5-2 and rs5-2 rs6-1,* under LD conditions. The size of the populations analyzed, from left to right, is n= 20, 20, 19, 18, 19, 18,19, 24, 24, 22, 17, 20 and 20. **B-C.** Flowering time of Col-0, *rs5-2* and *rs5-3* in LD conditions, measure as total leaf number (B) and days to bolting (C) (n= 17, 23 and 22, respectively). **D-E.** Flowering time of Col-0, *rs5-2* and *rs5-3* in LD conditions, measure as total leaf number (D) and days to bolting (E) (n= 31, 28 and 25, respectively). **** p-value < 0.0001; *** p-value < 0.001; ** p-value < 0.01; * p-value < 0.05

**Table 2.** Flowering time of Col-0 and simple and double mutants affecting raffinose metabolism grown in LD conditions.

| Genotype | Total leaf number |
| --- | --- |
| Col 0 | 15.1 ± 1.6 |
| *gols1-1* | 14.5 ± 1.8 |
| *gols2-1* | 14.5 ± 1.2 |
| *gols1-1 gols2-1* | 13.7 ± 1.7 |
| *gols3-1* | 14.8 ± 1.3 |
| *gols4-1* | 14.5 ± 1.2 |
| *rs2-1* | 14.9 ± 1.6 |
| *rs4-1* | 15.3 ± 1.6 |
| *rs5-2* | **12.2 ± 1.4 \*\*** |
| *rs5-3* | **13.5 ± 1.4 \*** |
| *rs6-1* | 15.0 ± 2.0 |
| *rs2-1 rs5-2* | **13.1 ± 1.3 \*\*** |
| *rs4-1 rs5-2* | **13.0 ± 1.1 \*\*** |
| *rs5-2 rs6-1* | **12.7 ± 1.1 \*\*** |

Since in our system, floral induction was triggered by activating the CO-FT module, we evaluated flowering time of the *rs5* mutants both in LD and SD conditions (Figure 5B-E). We confirmed that in LD conditions both mutant alleles flowered earlier than the wild type (Figure 5B-C). However, under SD conditions, only the *rs5-2* flowered significatively earlier than the wild type (Figure 5D-E). In addition, we observed that fertility of the *rs5-2* mutant was decreased, with a severe reduction in fruits per plant and seeds per silique (Figure 6A-B). The *rs5-3* mutant only displayed a slight reduction in the number of seeds per silique, and no alteration of the number of fruits (Figure 6A-B). Raffinose was not detected in leaves of the *rs5-2* mutant and moreover only residual raffinose synthase activity was present in leaf extract of this mutant and this activity did not increase in response to abiotic stress, as described for the wild type (Egert et al., 2013). Since biochemical characterization was more complete for the *rs5-2* mutant and we confirmed that it was a knock-out mutant, we decided to use this allele in further studies.

**Figure 6.**
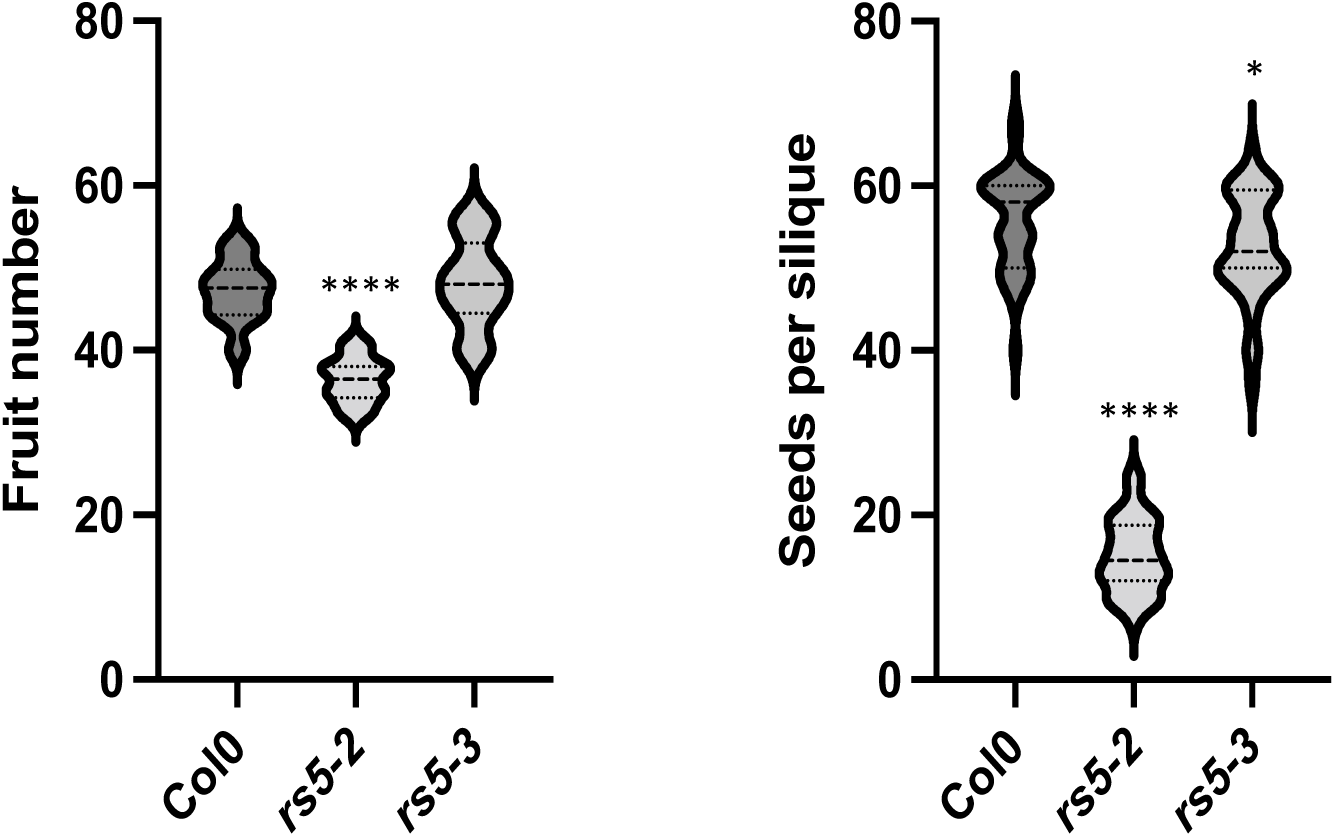
Characterization fruit number and seeds per silique in *rs5-2* and *rs5-3* mutants. **A.** Total fruit number produced by Col-0, *rs5-2* and *rs5-3* plants, scored when the shoot apical meristem arrested growth (n= 14). **B.** Seed number per silique in Col-0, *rs5-2* and *rs5-3* plants. The number of seeds was scores in ten siliques corresponding to nodes 6-15 for each plant. Vales represent the average ± SD of four biological replicates in F and G. ANOVA with Tukey correction was performed to calculate the significance differences. ***** p-value < 0.0001; * p-value < 0.05

### Expression of GOLS and RS genes in vegetative and reproductive developmental phases

As the results of the metabolomic analysis, the early flowering phenotype displayed by the rs5 mutants suggested as well that raffinose metabolism could play a role during floral transition. We therefore investigated the expression of *GOLS* and *RS* genes at different stages of development, analyzing samples of vegetative apices, apices undergoing floral transition and inflorescence apices (Figure 7). Upregulation of *SOC1* was used as a control indicating the initiation of floral transition. In agreement with observed changes in the transcriptome, expression of *GOLS1*, *GOLS3*, *RS2* and *GOLS4* decreased as plants switched from vegetative to reproductive stage. In addition, we found that *RS5* expression clearly decreased, meanwhile *RS6* expression temporarily increased during floral transition to decrease later in inflorescence apices. These observations, together with the early flowering phenotype of the *rs5-2 and rs5-3* mutants, points out to *RS5* as a putative candidate contributing to the control of floral transition via the regulation of the balance between simple and storage carbohydrates, such as raffinose, in the shoot apical meristem.

**Figure 7.**
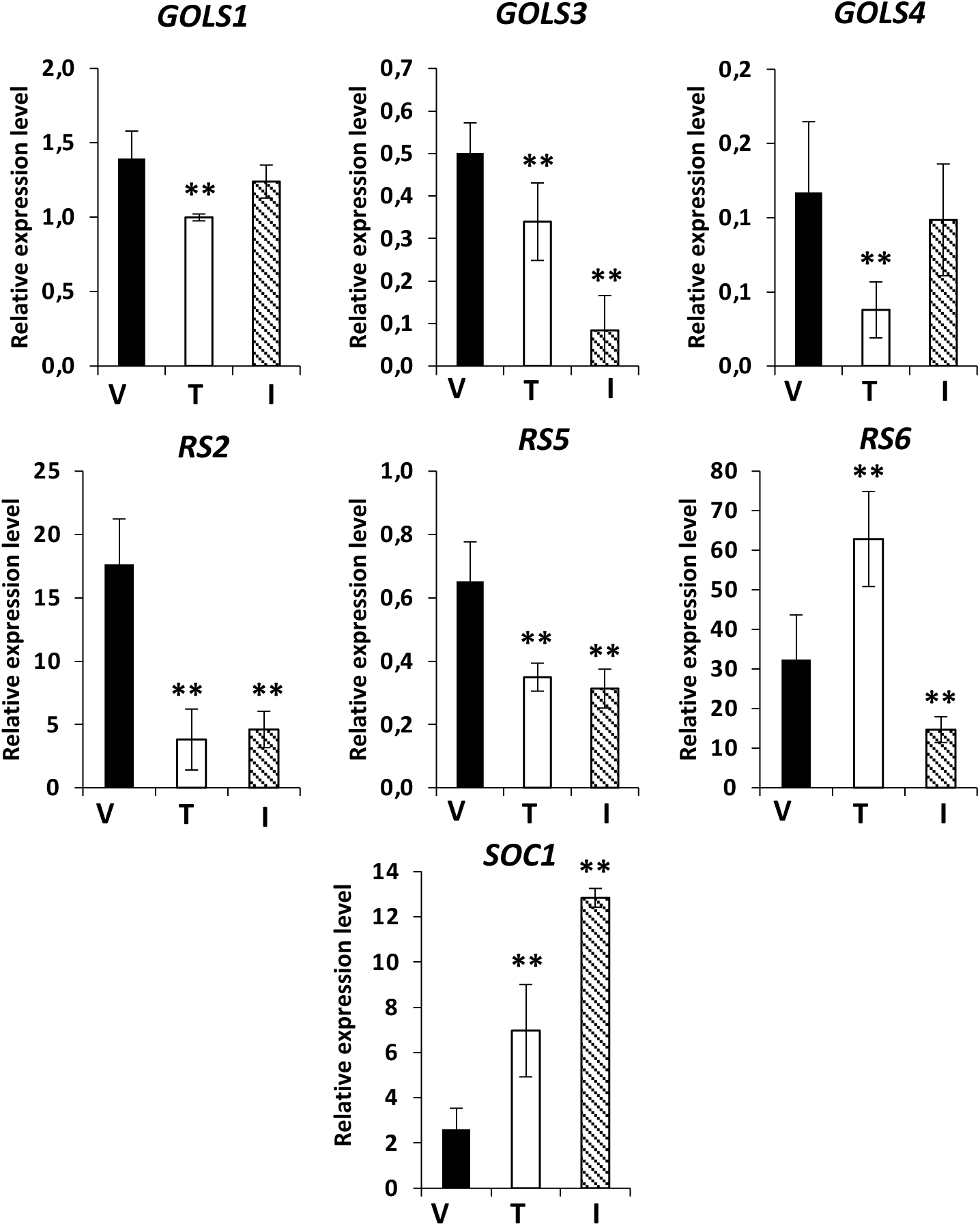
Relative expression level of genes involved in raffinose biosynthesis in vegetative, transition and inflorescence apices. *SOC1* expression was scored as a marker for floral transition. *TIP41* was used as a reference gene. V= vegetative apices on day 7. T= apices in floral transition on day 15. I= inflorescence apices on day 30. Results show the average ± SD of 3 biological replicates. Significance level measured by Anova with Tukey correction compared with the expression level on day 7 (vegetative stage). ** indicates p-value<0.01.

To further detail the spatial expression pattern of *RS5* in the meristem during phase transition we performed RNA *in situ* hybridization to analyze expression of *RS5* before, during and after floral transition (Figure 8). We detected strong *RS5* expression, at day 10 after germination, in the vegetative shoot apical meristem as well as in young leaf primordia (Figure 8A-B). Later in development, at day 12, *RS5* expression was less evident in the meristem but still patent in the leaf primordia (Figure 8C-D). Finally, at day 15, when floral transition has already taken place, there is almost no *RS5* expression in the inflorescence meristem and residual levels can be observed in older leaves (Figure 8E-F). This data confirmed that *RS5* is expressed in the meristem and the young leaf primordia before floral transition, as shown by the qRT-PCR data (Figure 7). This expression tends to decrease along with phase change, which would also agree with a difference in raffinose abundance found in the metabolomic data (Figure 2) and with the raffinose metabolism being pointed out as one of the pathways altered during floral transition (Table 1). With the aim of revealing the mechanism through which the raffinose metabolism impacted on the determination of floral transition, we investigated expression of flowering related genes in the *rs5-2* mutant.

**Figure 8.**
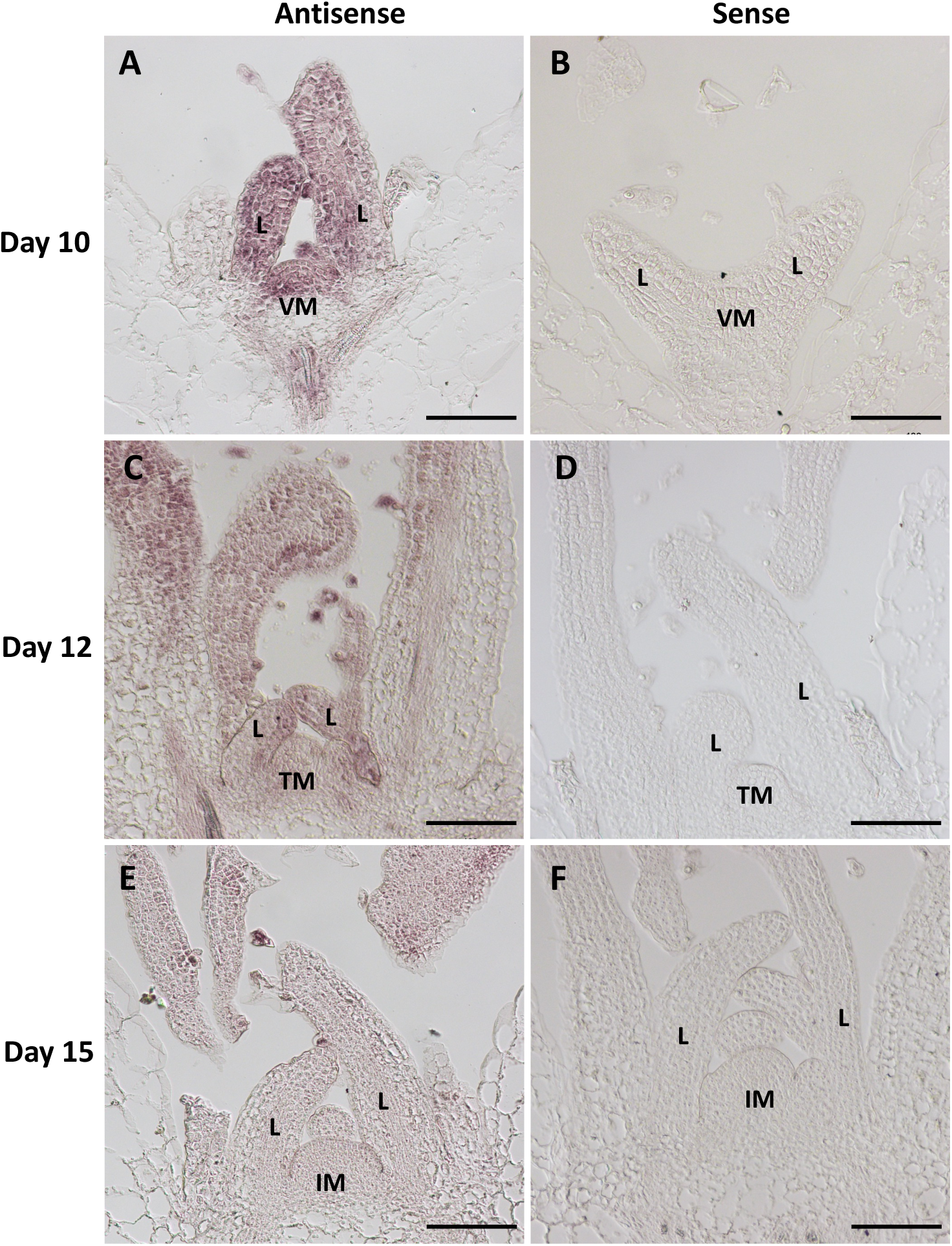
*RS5* expression pattern in the shoot apical meristem at different developmental stages. **A-B.** RNA *in situ* hybridization with *RS5* probe on shoot apices ten days after germination (vegetative stage) (A, antisense; B, sense). **C-D.** RNA *in situ* hybridization with *RS5* probe on shoot apices twelve days after germination (floral transition) (C, antisense; D, sense). **E-F.** RNA *in situ* hybridization with *RS5* probe on shoot apices twelve days after germination (inflorescence apex) (E, antisense; F, sense). VM = vegetative meristem; L = leaf primordia; TM = meristem in floral transition; IM = inflorescence meristem. Scale bars = 100 µm.

### Molecular characterization of the rs5-2 mutant

We characterized the expression of *SOC1*, *AP1*, *LFY* and *FT* in the *rs5-2* mutant compared to the wild type. To do so, we collected samples in a time course experiment with *in vitro-*grown plants, collecting samples (entire seedlings) every two days, between ZT14 and ZT16 (Figure 9). Expression of *AP1*, *SOC1* and *LFY* increased earlier in *rs5-2* mutant than in wild type seedlings, in agreement with the early flowering phenotype displayed by the mutant (Figure 9A-C). We could also observe that expression of the florigen *FT* was higher in *rs5-2* in all analyzed timepoints (Figure 9D). To determine whether it was only the onset of *FT* expression that was affected or also its diurnal expression, we performed a time-course experiment to compare *CO* and *FT* expression during the day (Figure 9E-F). We found that *CO* expression was very similar in both genotypes, with only minor differences at ZT10, ZT14 and ZT18 (Figure 9E). However, *FT* diurnal expression was significantly affected (Figure 9F). Interestingly, we found no significant differences in *FT* expression during the morning (ZT0-ZT6), but we did find an increased *FT* expression in *rs5-2* from ZT8 until ZT24, with a very marked difference close to the end of the day between ZT14 and ZT16.

**Figure 9.**
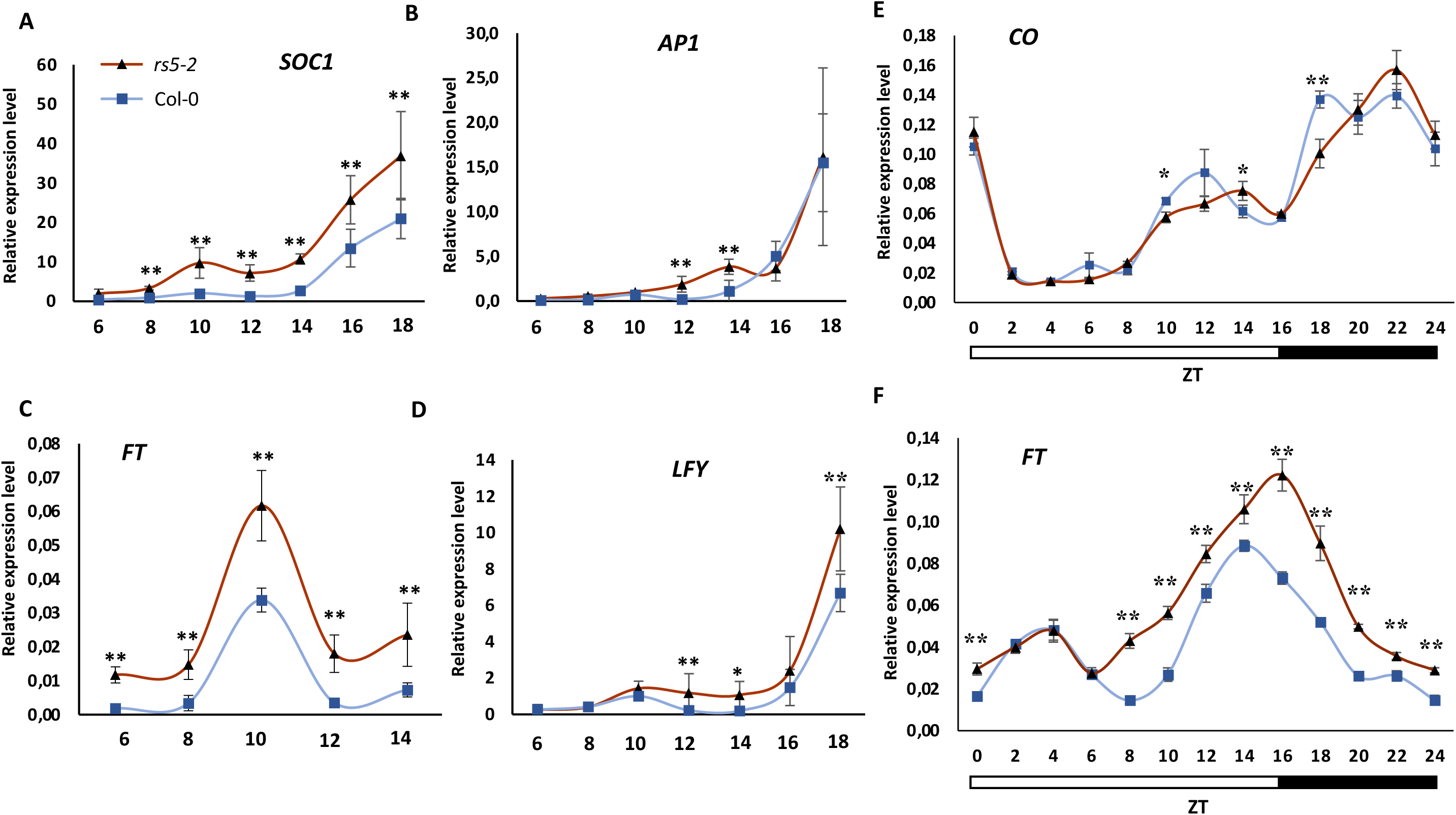
Expression of flowering genes, flowering time and floral meristem identity genes, in Col-0 and *rs5-2* mutant seedlings in LD. **A-D.** Time-course expression of *SOC1*, *AP1*, *FT* and *LFY* from day 6 until day 14 after germination (for FT) or 18 after germination (for the rest). Entire seedlings were collected every two days between ZT14 and ZT16. **E-F.** Time-course expression of *CO* and *FT* in LD. Entire seedlings were collected every two hours. *TIP41* gene was used as reference gene. Graphs represent the average ± SD of three biological replicates. Statistical significance was scored by Fisher t-test (* p-value<0.05, ** p-value<0.01).

To investigate how the defect in raffinose biosynthesis caused an early upregulation of the florigen and a consequent early floral transition in the *rs5-2* mutant, we measure the expression level of *THEHALOSE-6-PHOSPHATE SYNTHASE 1* (*TPS1*) and S*QUAMOSA PROMOTER BINDING PROTEIN-LIKE 3* (*SPL3*), two genes encoding key proteins in the carbohydrate status sensing mechanism in Arabidopsis (Figure 10A-B). The results show that *TPS1* expression is increased in the *rs5-2* mutant compared to the wild type, as soon as six days after germination, and displays higher expression at all time points tested (Figure 10A). It has been shown that an increase in TPS1 activity leads to increased T6P levels that promotes *SPL3* expression (Wahl et al., 2013). In agreement with that, we also observed that *SPL3* displayed a higher expression at day 10 in the *rs5-2* mutant (Figure 10B). We have previously described a decrease in raffinose on day 1 and an increase in mono and disaccharides on day 3 (Supplementary Figure 3). These changes impact on *TPS1* expression, probably increasing the levels of trehalose-6-phosphate, and therefore impact on the timing of floral transition.

**Figure 10.**
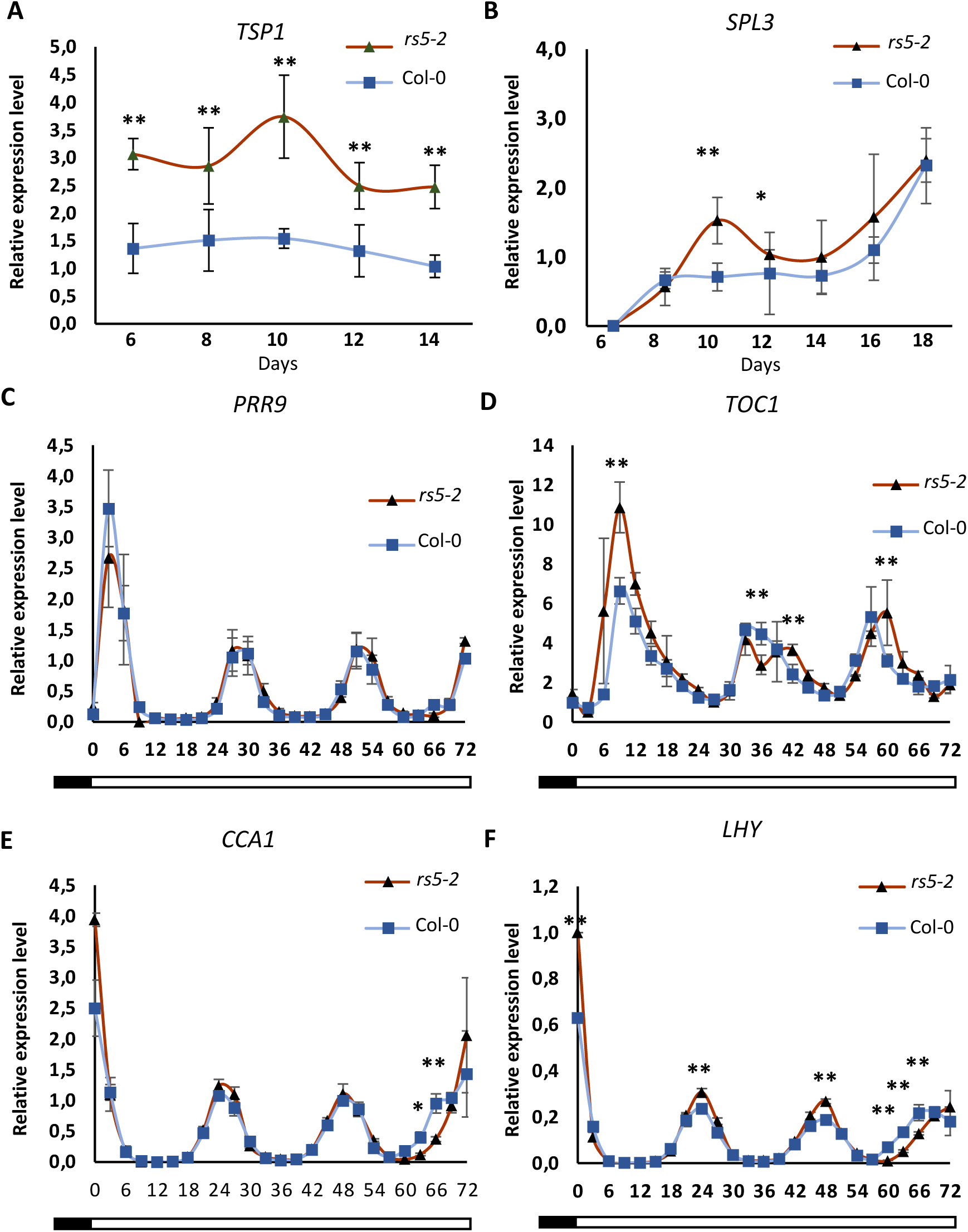
Expression of genes related to trehalose synthesis and signaling and circadian clock function in Col-0 and *rs5-2* mutant seedlings. **A-B.** Time-course expression of *TPS1* and *SPL3* from day 6 until day 14 after germination (TPS1) or 18 after germination (*SPL3*). Entire seedlings were collected every two days between ZT14 and ZT16. *TIP41* gene was used as reference gene. Graphs represent the average ± SD of three biological replicates. **C-F.** Time-course expression of *PRR9, TOC1, CCA1* and *LHY* in free-running conditions. Seedling were entrained for ten days in 12/12 dark/light cycles before being transferred to continuous light. 16 seedlings per biological replicate were collected every 3 hours for 3 days. *IPP2* gene was used as reference gene. Graphs represent the average ± SD of three biological replicates

Since carbohydrate status has been shown to influence the circadian clock (Haydon et al., 2017), we investigated whether expression of core clock oscillator genes and related regulatory loops are altered in the *rs5-2* mutant. With this aim, we grew Col-0 and *rs5-2* plants under day-neutral conditions for 10 days (12 h light, 12 h dark) and then we switched them to free-running conditions (continuous light) for 3 days and collected samples (entire seedlings) every 3 hours. We quantified expression of core circadian clock genes, among them the morning genes *CIRCADIAN CLOCK-ASSOCIATED 1* (*CCA1*) and *LATE ELONGATED HYPOCOTYL* (*LHY*), the evening gene *TIMING OF CAB EXPRESSION 1* (*TOC1*) and *PSEUDO-RESPONSE REGULATOR 9* (*PRR9*), a member of the so-called regulatory morning-loop (Figure 10C-F). There were no major changes in the expression of the core clock genes, although we could observe a delay in *CCA1*, *LHY* and *TOC1* upregulation after three days in free-running conditions. Regarding *PRR9* expression, we could not detect any significant difference between Col-0 and *rs5-2*. These preliminary results suggest a long period phenotype, however further confirmation would be needed to validate the effect of the *rs5-2* mutant in the clock, and therefore the potential contribution of this effect on the early flowering phenotype of the mutant.

Finally, since our results point out that a different balance between raffinose and simple carbohydrates leads to an alteration of flowering time, we decided to compare the abundance of different carbohydrates in apices and leaves of the *rs5-2* mutant in comparison to Col-0 plants. We collected samples on day 12 after sowing, when according to our data the *rs5-2* mutant already exhibits *AP1* expression (meaning that floral transition has already occurred on those plants). Col-0 plants on day 12 do not display *AP1* expression, which becomes evident at day 14-16, meaning that these plants are undergoing floral transition. We observed major changes in carbohydrate abundance in both leaf and apex samples (Table 3). We confirmed that raffinose levels were significantly lower in the mutant, although this difference was only apparent in leaf samples and not in apex samples. According to the decrease in raffinose, we observed a major increase in galactinol both in the leaves and in the apices. Moreover, we detected an increase in amounts of other carbohydrates such as erythritol and fructose in *rs5-2* mutant. On the contrary, rhamnose, glucose, maltose and myo-inositol showed lower abundance in the *rs5-2* mutant than in Col-0.

**Table 3.**
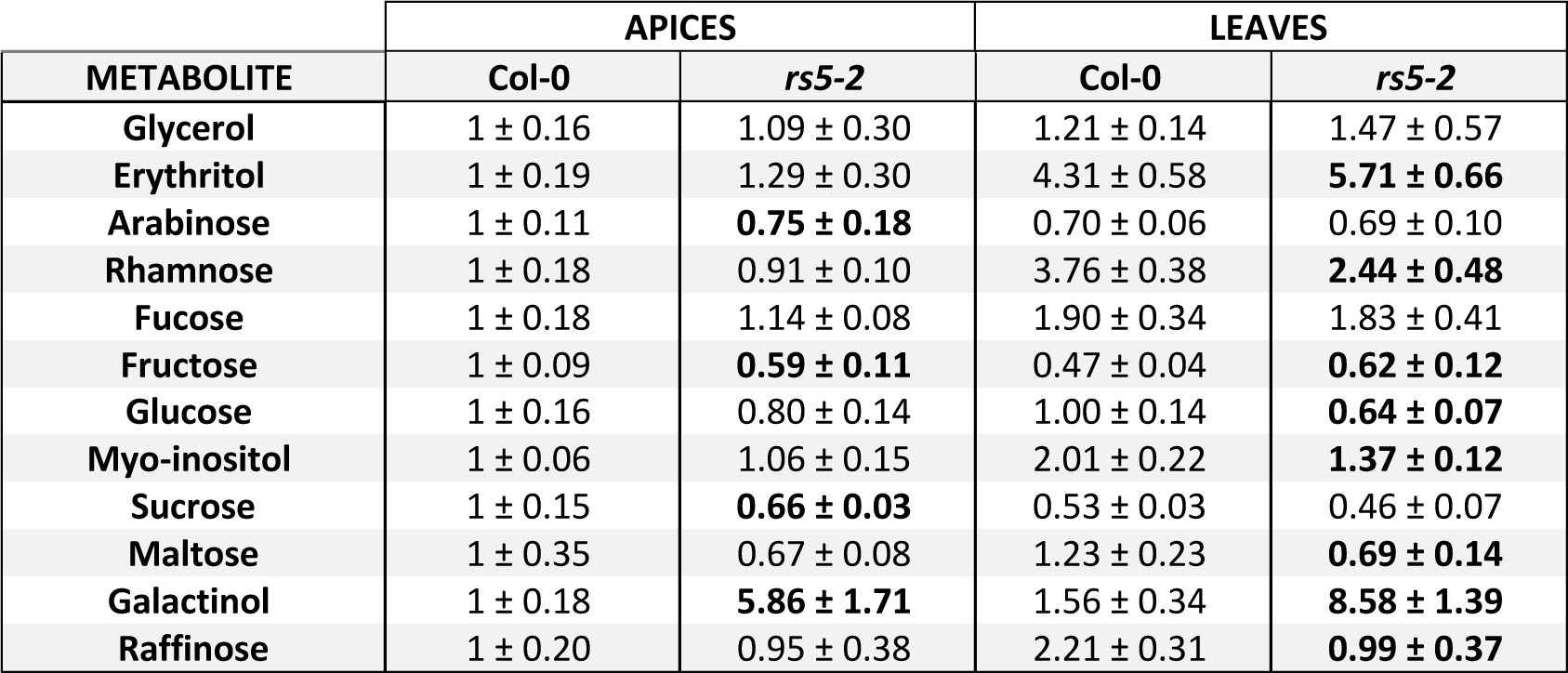
Carbohydrates detected by GC-MS in apex and leaf samples from the *rs5-2* mutant and Col-0.

In summary, this work provides new insights on the metabolic changes that accompany floral transition at the shoot apical meristem in Arabidopsis. Several pathways have been pointed out and will be the subject for further research. Our data suggest that changes in raffinose metabolism are relevant during the transition from vegetative to inflorescence meristem. We have identified differences in raffinose and mono/disaccharides accumulation in vegetative and inflorescence shoot apices and we have investigated the molecular consequences of a loss-of-function in one of the major raffinose synthase enzymes. Alteration of the raffinose synthesis leads to early flowering and altered expression of genes related to trehalose synthesis and signaling. These data suggest that raffinose acts as carbohydrate pool that is metabolized to simple sugars and used during floral transition, to produce reproductive structures.

## Discussion

In this work, we present a detailed characterization of the metabolic changes associated with floral transition in Arabidopsis. We have generated a dexamethasone inducible system based on the expression of a CO::GR fusion protein under the control of *CO* endogenous promoter. We have shown that a 3 kb CO promoter replicates that of the endogenous *CO* gene, both in spatial and temporal expression, and therefore dives the expression of the *CO::GR* transgene to those cells that normally would express endogenous *CO*. Upon dexamethasone treatment, CO-GR protein triggers floral induction by activating *FT* expression, mimicking the molecular events taking place in the leaves during photoperiod-dependent floral induction by the endogenous CO-FT module. This inducible system is robust, as shown by the complementation of the late-flowering phenotype in dexa-treated pCO::CO::GR *co-10* plants compared to mock-treated plants of the same line. Similar systems have been previously used to characterize the architecture of the genetic network responsible for triggering floral transition downstream of CO (An et al., 2004; Simon et al., 1996).

Several studies have identified relevant metabolites that accumulate or decrease during reproductive phase change (Conti, 2017; Gawarecka & Ahn, 2021; Olas et al., 2021). However, a global characterization integrating transcriptomic and metabolomic changes along the floral induction process was still lacking. Our data shows that the process of floral induction encompasses a metabolic reprogramming affecting both primary and secondary metabolism. In Arabidopsis plants induced to flower via a change in photoperiod (SD-to-LD shift) or via the activation of the CO-FT module in the leaf, there is a time gap that varies from 1 to 5 days, depending on the system and the conditions, until there is a clear morphological change in the shoot apical meristem (Abe et al., 2005; Simon et al., 1996; You et al., 2017). This change is characterized by the doming of the meristem and the formation of floral meristems at its flanks. It is logical to think that such morphological adjustment must be preceded by changes in relevant metabolites. Our data set show that there are very few changes in the leaf in response to the activation of CO. This agrees with the few changes we found in the leaves at the transcriptional level, including upregulation of *FT* on day one after the dexamethasone treatment. In the apex, on the contrary, there are major changes in metabolites that point out to several pathways as being significantly altered during photoperiod-dependent floral transition. Global analysis of metabolite changes in the apex allowed us to differentiate between an “early transition” phase, corresponding to day 1 after induction, and a “late transition” phase corresponding to day 3. Expression of floral markers, *LFY* and *AP1*, on day 5 evidence that floral transition has been accomplished by this time. Metabolic changes on the early transition affect both carbon and nitrogen metabolism: there is a marked decrease in several amino acids, such as phenylalanine, tryptophan, proline, glutamine, threonine, valine, asparagine and alanine, and a decrease in raffinose, a major storage carbohydrate. Along with this, myo-inositol decreases and galacturonic acid, an oxidized form of galactose, accumulates. These three metabolites, raffinose, myoinositol and galactose, are involved in RFOs biosynthesis. Changes in amino acids during floral transition has been previously observed in Arabidopsis (Corbesier et al., 2001) and more recently specific amino acid signatures have been proposed for vegetative and reproductive stages (Olas et al., 2021).

On the other hand, the late transition phase, between day 3 and day 5 after treatment, is characterized by an increase in mono and disaccharides, such as fructose, galactose, arabinose, glucose and sucrose, among others. This increase in simple carbohydrates precedes the morphological changes that occurs in the shoot apical meristem once floral transition occurs. Changes in gene expression confirm that raffinose metabolism, or RFOs biosynthesis, is one of the pathways that is significantly altered during floral transition. Among genes involved in raffinose synthesis, several galactinol synthase genes (*GOLS1*, *3* and *4*) and one raffinose synthase gene, *RS2*, decrease their expression during floral transition.

*GOLS* genes catalyze the first step in RFO biosynthesis, that is the synthesis of galactinol from UDP-galactose and myo-inositol. In Arabidopsis there is seven members of the *GOLS* family (*GOLS1*-*GOLS7*) and *GOLS1*, *2* and *3* contribute to the plant response to oxidative stress and drought tolerance, both in Arabidopsis and chickpea(Nishizawa et al., 2008; Salvi et al., 2018; C. Song et al., 2016; Taji et al., 2002). *GOLS* genes have been also shown to regulate seed germination, longevity and vigor (Jang et al., 2018; Salvi et al., 2016). Interestingly, *MdGOLS1* is differentially expressed during bud dormancy and bud break in apple. The accumulation of RFOs in dormant buds and their decrease prior to bud break suggest that these metabolites display a dual role offering protection against dehydration (during bud dormancy) and source of energy when buds get reactivated (da Silveira Falavigna et al., 2018). *RS* genes, on the other hand, catalyze the synthesis of raffinose from galactinol and galactose. Six genes have been predicted to act as raffinose synthases in Arabidopsis (*RS1*-*RS6*), however raffinose synthase activity has only been biochemically demonstrated for RS4 and RS5 (Nishizawa et al., 2008). In fact, RS2 acts as an alpha-galactosidase (degrading raffinose in sink organs), RS4 acts mainly as a high affinity stachyose synthase, although it also displays raffinose synthase activity and galactosyl hydrolase (degrading galactinol and stachyose) (Gangl et al., 2015; Peters et al., 2010). *RS6* expression is induced under drought stress and cold conditions and contributes to the synthesis of raffinose. Finally, RS1 has not been functionally characterized and *RS3* is a pseudogene (Gangl & Tenhaken, 2016). Therefore, RS5 is the major specific raffinose synthase in Arabidopsis (Egert et al., 2013). Taken this into account and given the expression changes of *RS5* in the apex during floral transition as shown by RNA *in situ* hybridization, we focused our efforts on the characterization of the consequences that a loss-of-function mutation on the *RS5* gene has on flowering time.

Several pieces of information have previous related raffinose metabolism to the control of flowering time. Mateos and colleagues (2011) showed that the *din10* mutant, a knock-out affecting *DARK INDUCED 10*/*RS6* gene, displays a slight early flowering phenotype with an early and increased *FT* expression. *RS6* expression in response to cold exposure is regulated by the complex FLOWERING LOCUS C (FLC)-SHORT VEGETATIVE PHASE (SVP) (Mateos et al., 2011). Besides, heterologous analysis of the expression pattern of the *Cucumis melo GALACTINOL SYNTHASE 1* (*CmGAS1)* gene showed that its promoter was active in a subset of phloem companion cell in minor veins of Arabidopsis and tobacco, in the exact same cells that activate expression of *FT* in response to the photoperiod (Chen et al., 2018; Haritatos et al., 2000). In fact, ablation of the cells in which the *CmGAS1* promoter is active delays flowering in tobacco (Chen et al., 2018). In this line, overexpression of Arabidopsis *GOLS2* gene in rice confers drought tolerance under field conditions, increasing grain yield with an associated early flowering phenotype (Gomez Selvaraj et al., 2017).

In this context, our work shows that a deficiency of raffinose synthesis, due to a loss-of-function mutation affecting the *RS5* gene, causes an early flowering phenotype. We analyzed two mutant alleles, rs5-2 and *rs5-3*, both showing an early flowering phenotype in LD and defects on fertility. Phenotype of the *rs5-2* was more severe in all aspects than that of the *rs5-3* allele. To identify if there was redundancy among *RS* genes, we analyze mutants affecting *RS2*, *RS4* and *RS6*. We could not confirm that *RS6* loss-of-function causes an early flowering phenotype, as opposed to what it was described for the *din10* mutant, since under our growth conditions the *rs6-1* mutant (SALK035336 line) did not show any flowering phenotype. Since the early flowering phenotype displayed by the *din10* mutant was not mild, this discrepancy could be due to slight differences in growth conditions.

An alteration in the balance between accumulation of storage carbohydrates, such as raffinose, and the release of mono and disaccharides, seems to be a relevant change occurring during floral transition, prior to the upregulation of floral marker genes and morphological changes at the shoot apical meristem. This alteration of the ratio between storage oligosaccharides and simple carbohydrates suggests that a metabolic reprogramming occurs during floral transition, in agreement with a higher energy demand required to generate reproductive tissues (sink tissues), flowers and fruits, as compared to generation of leaves (source tissues) during the vegetative development. Plants are characterized by their developmental plasticity and reproductive phase change corresponds to one of the major trade-offs between survival, growth and reproduction (Gnan et al., 2017; Lundgren & des Marais, 2020). Our results suggest that a change in the ratio between mono/disaccharides and raffinose in the apex could play a role in determining flowering time. A similar idea has been proposed for the control of seed vigor by RFOs in Arabidopsis and maize: a high RFO/sucrose ratio correlates with increased seed vigor, and a sharp decrease in raffinose levels have been observed during seed imbibition and early germination in several legume species (T. Li et al., 2017; Salvi et al., 2018; Vidal-Valverde et al., 2002). Also, a decrease in raffinose is observed in apple trees prior to bud break, corresponding to a lower raffinose/sucrose ratio at the time when the bud becomes metabolically active (da Silveira Falavigna et al., 2018). A similar result was observed in chestnut trees, in which induction of *GOLS* genes is related to winter endodormancy (Ibáñez et al., 2013). We here propose that the reproductive phase change is correlated in Arabidopsis with a change in the ratio between raffinose and mono/disaccharides, with a lower ratio corresponding to a meristem undergoing floral transition in comparison to a vegetative meristem (Figure 11).

**Figure 11.**
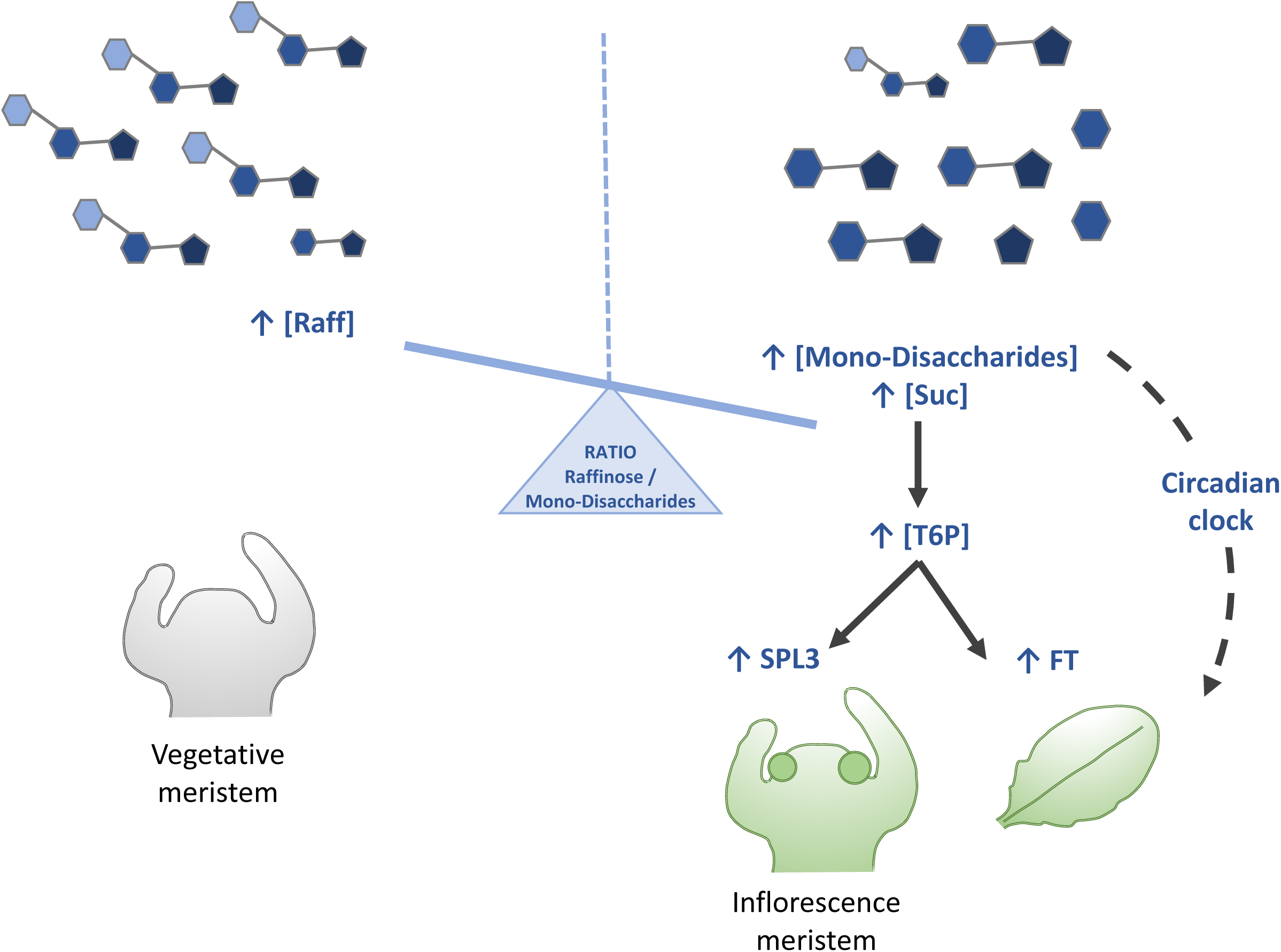
Model on how raffinose/mono-disaccharide ratio can influence flowering time. Raffinose accumulates in the shoot apical meristem during vegetative development. Upon perception of the photoperiod signal, raffinose synthesis decreases and the amount of mono- and disaccharides increase. A lower raff/mono-disaccharides in the shoot apex contributes to the floral transition. Increase sucrose correlates increased TPS1 expression and accumulation of T6P. T6P in turn exerts its function through upregulation of FT in the leaf and SPL3 in the apex, both changes accelerating floral transition. On the other hand, increased sucrose affects the function of the circadian clock, among others stabilizing the FT-activator GI.

We investigated how changes in the raffinose/mono-disaccharides ratio could influence flowering time. In Arabidopsis, the trehalose-6-phosphate (T6P) pathway integrates the carbohydrate status and contributes to the fine-tuning of the optimal time for flowering. In plants, trehalose is synthesized from glucose-6-phosphate and UDP-glucose via the intermediate T6P. *TREHALOSE PHOSPHATE SYNTHASE1* (*TPS1*) catalyzes the production of T6P, which is subsequently dephosphorylated to trehalose by *TREHALOSE PHOSPHATE PHOSPHATASES* (*TPP*s) (Paul et al., 2008). T6P acts as a signal of sugar availability and modulates both vegetative and reproductive phase change (Ponnu et al., 2020; Wahl et al., 2013). In leaves, T6P level shows a diurnal fluctuation very similar to that of *FT* expression, with a maximum at the end of the light period in long days. Wahl et al. showed that T6P is needed for the upregulation of *FT* expression in phloem companion cells. In addition, T6P also regulates *SQUAMOSA PROMOTER BINDING PROTEIN-LIKE* (*SPLs*) genes in the SAM via miRNA156-dependent and independent pathways (Wahl et al., 2013). In this context, we hypothesize that a decrease in the raffinose/mono-disaccharides ratio in the shoot apical meristem could cause an upregulation of *TPS1* and a concomitant increment of T6P. Then, increased T6P levels contribute to the early upregulation of *FT* in the leaves of the *rs5-2* mutant and to the upregulation of *SPL3* in the apex. These changes in gene expression explain the early flowering phenotype of the *rs5-2* mutant. Nevertheless, changes in *FT* gene expression could also be explained by the effect of an increased raffinose/mono-disaccharides ratio on the circadian clock function. Daily changes in photosynthesis-derived sucrose are important for circadian clock entrainment through repression of the morning-expressed gene *PSEUDO-RESPONSE REGULATOR 7* (*PRR7*). On the other hand, sucrose affects GIGANTEA stability, a known activator of *FT* (Haydon et al., 2017). GI affects photoperiod flowering in different ways: it controls CO stability, it direct binds *FT* promoter and interacts with FT-repressing factors such as SHORT VEGETATIVE PHASE (SVP), TEMPRANILLO 1 (TEM1) and TEMPRANILLO 2 (TEM2) (Sawa & Kay, 2011; Y. H. Song et al., 2014). Our experiments show a mild effect of the *rs5-2* mutation on the clock function, affecting the amplitude of *TOC1* expression and possibly the period of *TOC1*, *LHY* and *CCA1*. These changes in clock gene expression could also contribute to an altered output response, such an early *FT* upregulation.

Our data on quantification of sugar concentrations on the leaves and the apex of the *rs5-2* mutant compared to the wild type showed contradictory results, not in line with our proposed model inferred from changes identified though metabolomic and transcriptomic data. However, the *rs5-2* mutant is expected to show a deficiency in raffinose synthesis in all tissues where the *RS5* gene is expressed. These defects are probably more extensive and complex than the effect we could expect from changes in *RS5* gene expression in the wild type plant during photoperiodic activation of floral induction. In this way, we identified changes in raffinose content in leaves of the *rs5-2* mutant that we could not observed in dexamethasone *versus* mock-treated plants. On the contrary, we could not observe changes in raffinose content in the apex of the *rs5-1* mutant compared to the wt. This could be explained by the fact that sugar quantification was performed in 12-day old seedlings, a time point at which *rs5-2* has already undergone floral transition, while Col0 seedlings still are in a vegetative state. In addition, in our initial experimental model, we identified molecular events taking place in response to the activation of CO, which might differ to what occurs when plants are expose to natural long photoperiod.

In summary, our data suggest that a change in the raff/mono-disaccharides ratio occurs in the apex during floral induction. These changes are possibly integrated by the T6P pathway and the circadian clock function and contribute therefore to the fine-tuning of flowering time in Arabidopsis. It is possible that accumulation of raffinose plays a regulatory role in organs with different metabolic activity when they go through different developmental stages, such as seeds (dormant/germinating), buds (budset/budburst) or meristems (vegetative/reproductive). Tissue specific modification of raffinose levels, in wild type and mutants from flowering pathways, could help to confirm if this is the case in vegetative *versus* inflorescence meristems and to clarify the role of raffinose in the control of flowering.

## Supporting information

Supplementary Table 3. Identified metabolites in leaf and apex samples on day 0, 1, 3 and 5.

Supplementary Table 4. Metabolites with significant changes in the apex across the experiment.

Supplementary Table 5. Analysis of enriched pathways by Metaboanalyst and Plant Metabolic Network platforms in apex samples on day 1 and day 3.

Supplementary Table 6. Differential expressed genes in apex and leaf samples on day 1 and day 3.

## Acknowledgements

This work was supported by the MCIN/AEI /10.13039/501100011033 to RB (grants nos BIO2015-73491-JIN funded by MCIN/AEI and FEDER Una manera de hacer Europa and PID2020-113035RB-I00 by MCIN/AEI) and to FM (grants nos BIO2015-64307-R and PGC2018-099232-B-I00 by MCIN/AEI and FEDER Una manera de hacer Europa). JP was supported by a contract for personal predoctoral en formación (grant no BES-2016-078276 by MCIN/AEI y por FSE invierte en tu futuro). We thank the Metabolomics Service at IBMCP (UPV-CSIC) for technical support in the quantification of sugars. We thank the Swedish Metabolomic Centre for technical assistance and data processing and the UPSC Bioinformatics Facility for transcriptome data analysis. We thank Dr A Berbel for advice and help with *in situ* hybridization experiments and MJ Domenech for efficient technical assistance.

## Supplementary Figures

**Supplementary Figure 1.**
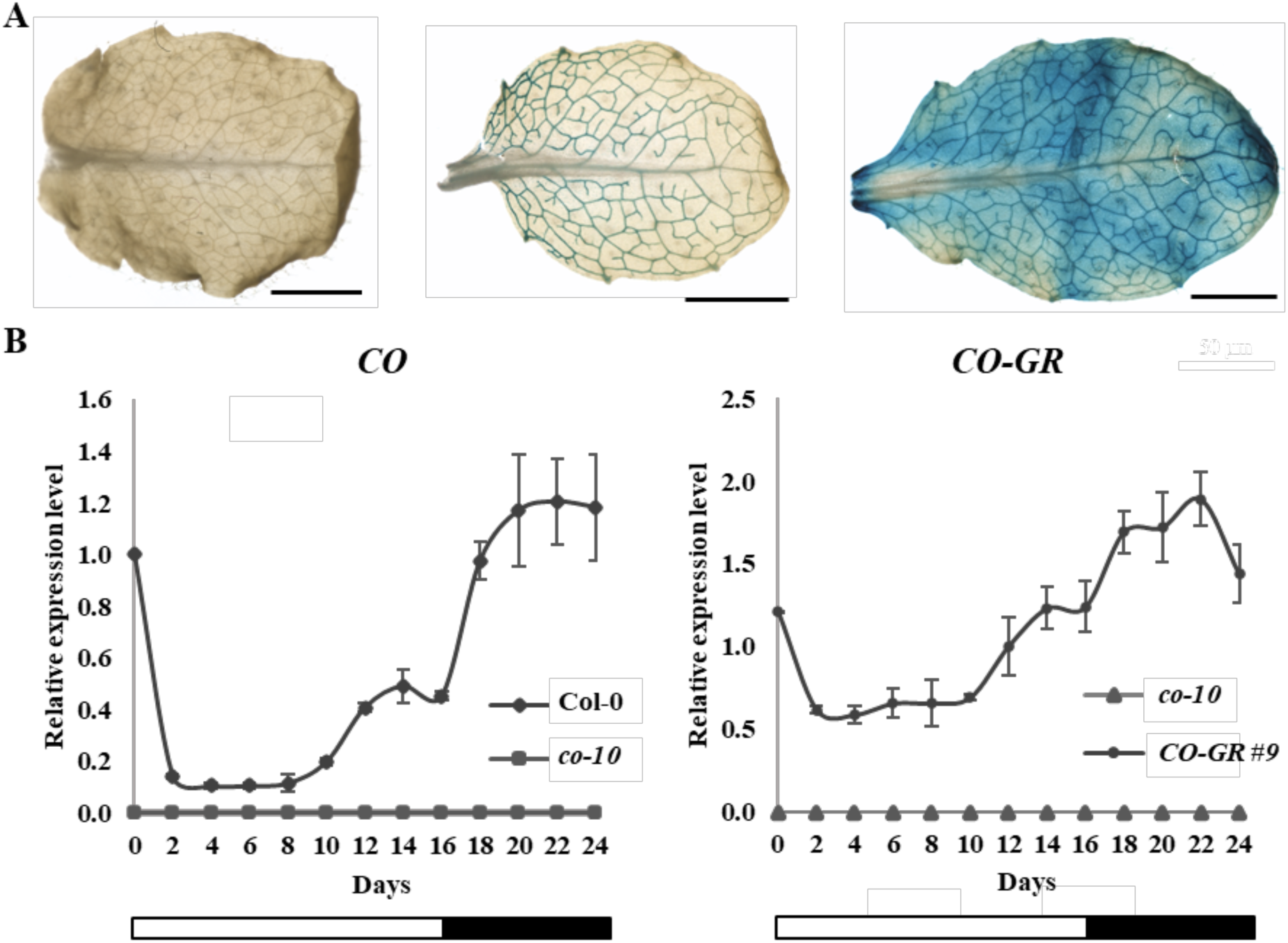
Characterization of the spatial and temporal expression pattern driven by the *CO* promoter (*pCO*) used in the inducible system. **A.** GUS assay in Col-0 (negative control), *pCO::GUS* transgenic line and *35S::GUS* (positive control). **B.** Time-course expression of the *CO-GR* transgene under the *CO* promoter in the selected p*CO*::*CO-GR co-10* #9 line. Left panel, *CO* expression in Col-0; right panel, *CO-GR* expression in the #9 line. Expression levels are the average of three biological replicates, error bars correspond to the SD. IPP2 was sued as a reference gene.

**Supplementary Figure 2.**
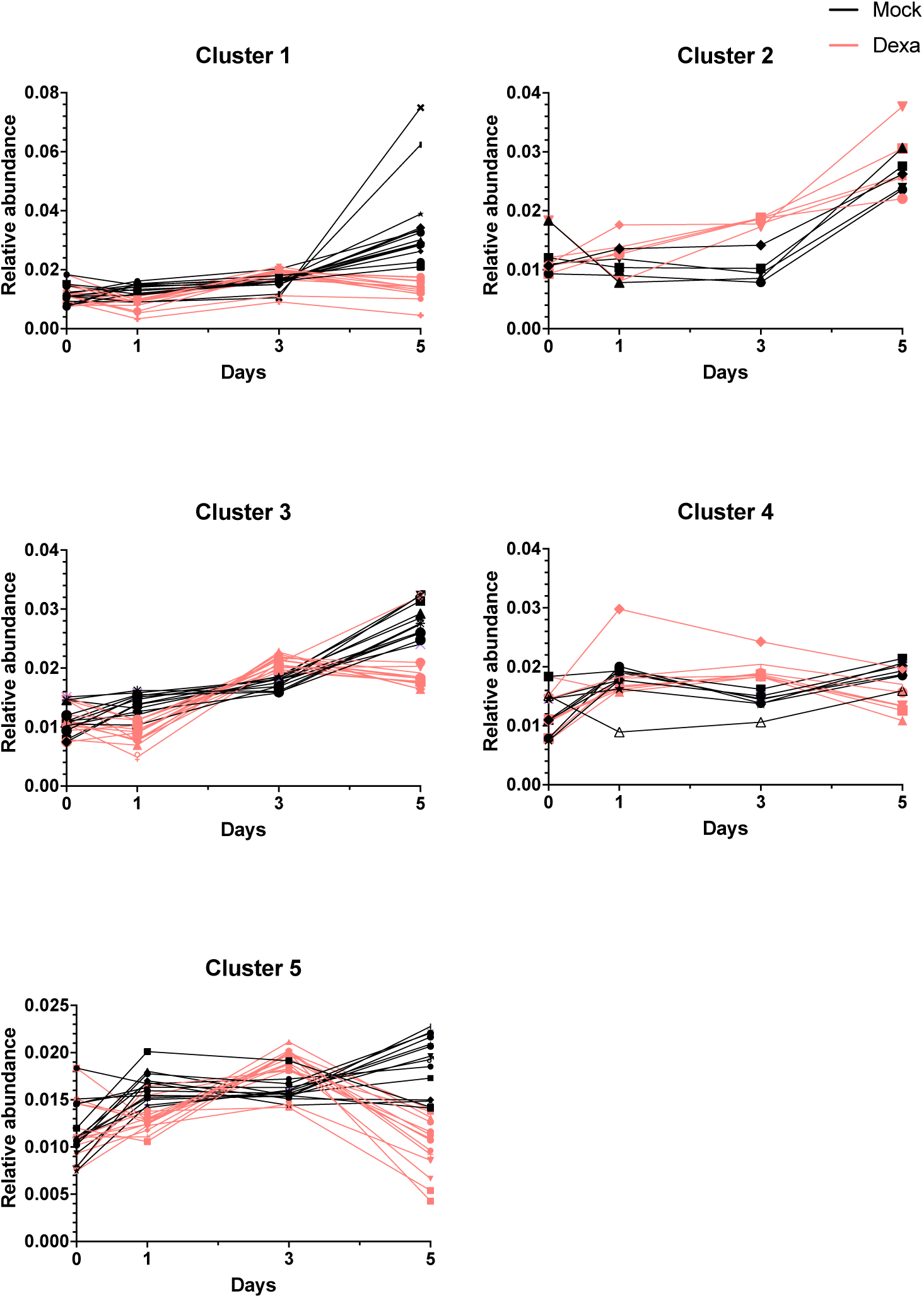
Hierarchical clustering of the top 55 metabolites with more significant changes across the experiment. Cluster 1 to cluster 5 correspond to the clusters identified in Figure 2.

**Supplementary Figure 3.**
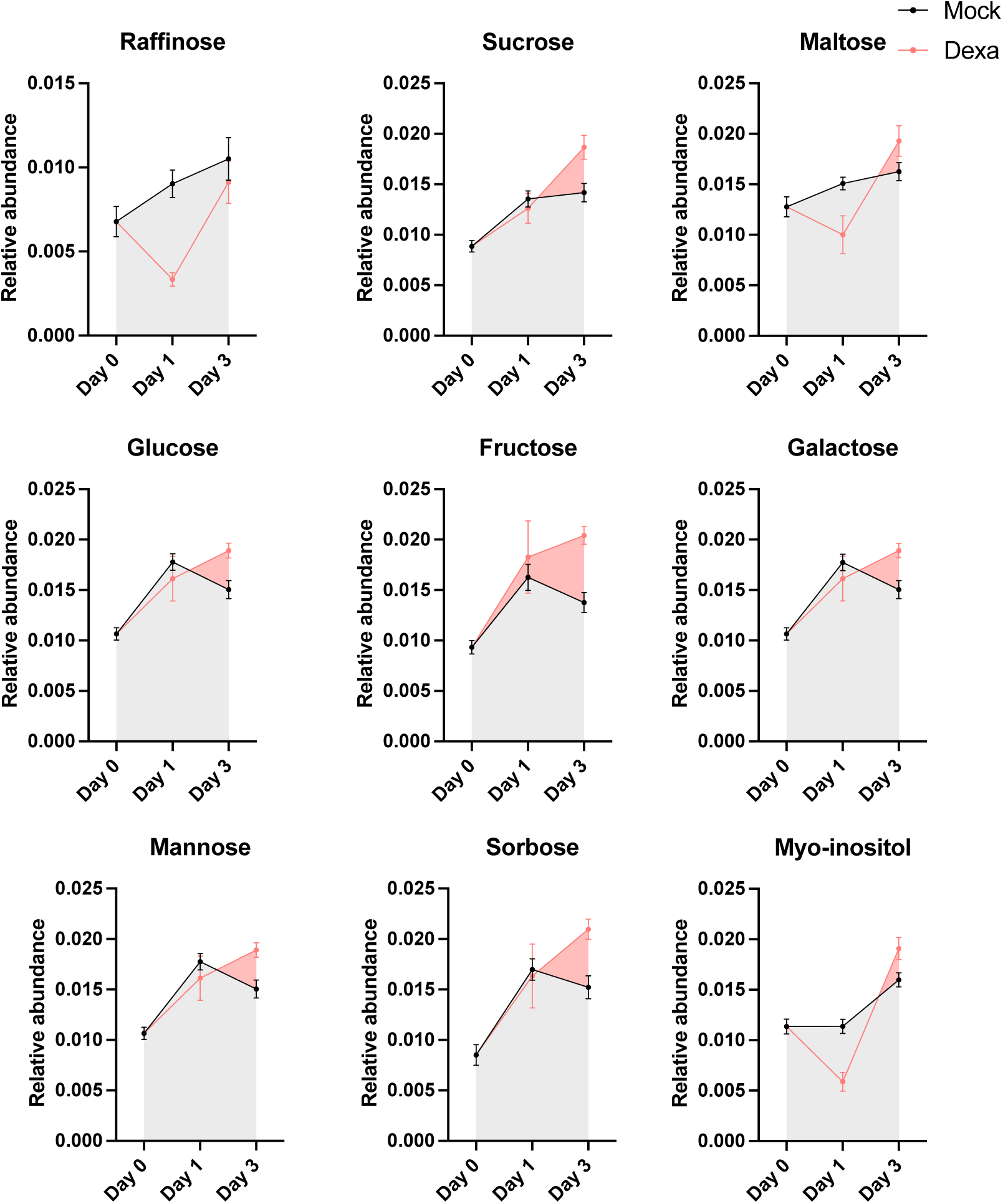
Changes in carbohydrates and myo-inositol on day 1 and 3 in dexa and mock-treated samples.

## Supplementary Tables

**Supplementary Table 1.** Flowering time response in dexamethasone-inducible transgenic lines in Col0, *ft-10* and *co-10* mutant backgrounds. Twelve independent T1 lines were analyzed for each construct and mutant background. Approximately 100 T2 seeds were grown in vitro in MS medium supplemented with basta to asses the number of insertions in each lines. Lines with one T-DNA copy were selected and, for each of them, 24 plants were acclimated in the greenhouse. One week after, plants were treated with desamethasone (30 uM dexamethasone 0.01% Tween-20) or mock (0.01% Tween-20). Flowering time was assessed as total leaf number and the result was standardized as compared to the corresponding background genotype: S (response similar to the background), LF (late flowering response) and EF (early flowering response).

| Background | Construct | Response |  |
| --- | --- | --- | --- |
|  |  | Dexamethasone | Mock |
| Col-0 | *pSUC2::CO-GR* | S | LF |
| Col-0 | *pSUC2::FT-GR* | S | S |
| Col-0 | *pFT::FT-GR* | S | S |
| Col-0 | *pCO::CO-GR* | S | LF |
| *ft-10* | *pSUC2::CO-GR* | S | S |
| *ft-10* | *pSUC2::FT-GR* | S | S |
| *ft-10* | *pFT::FT-GR* | S | S |
| *ft-10* | *pCO::CO-GR* | S | S |
| *co-10* | *pSUC2::CO-GR* | EF | S |
| *co-10* | *pSUC2::FT-GR* | S | S |
| *co-10* | *pFT::FT-GR* | S | S |
| *co-10* | *pCO::CO-GR* | EF | S |

**Supplementary Table 2.** Flowering time determination in homozygous transgenic lines pCO::CO-GR co-10 (T3 plants) upon dexamethasone or mock treatment. Data shows the total leaf number (average ±SD); n=25).

| Controls | Construction | Total leaf number |  |
| --- | --- | --- | --- |
| Col-0 | - | 18.4 ± 1.6 | |
| *co-10* | - | 46.1 ± 4.4 | |
| Line |  | Dexa | Mock |
| #2 | *pCO::CO-GR* | 29 ± 2.6 | 44.5 ± 3.6 |
| #3 | *pCO::CO-GR* | 33 ± 2.3 | 45.5 ± 3.7 |
| #4 | *pCO::CO-GR* | 26.2 ± 2.4 | 44 ± 3.3 |
| #7 | *pCO::CO-GR* | 29 ± 2.7 | 44.4 ± 4.3 |
| #9 | ***pCO::CO-GR*** | **25 ± 1.7** | **45.2 ± 3** |
| #1 | *pSUC2::CO-GR* | 28.6 ± 1.9 | 46.4 ± 3.8 |
| #2 | *pSUC2::CO-GR* | 35.4 ± 2.8 | 45.8 ± 2.5 |
| #3 | *pSUC2::CO-GR* | 33.5 ± 2.4 | 46.3 ± 3.4 |
| #5 | *pSUC2::CO-GR* | 35.9 ± 2.7 | 46.3 ± 2.7 |
| #8 | *pSUC2::CO-GR* | 32.4 ± 2.4 | 45.1 ± 3.5 |

**Supplementary Table 3.** Identified metabolites in leaf and apex samples on day 0, 1, 3 and 5.

**Supplementary Table 4.** Metabolites with significant changes in the apex across the experiment.

**Supplementary Table 5.** Analysis of enriched pathways by Metaboanalyst and Plant Metabolic Network platforms in apex samples on day 1 and day 3.

**Supplementary Table 6.** Differential expressed genes in apex and leaf samples on day 1 and day 3.

**Supplementary Table 7.**
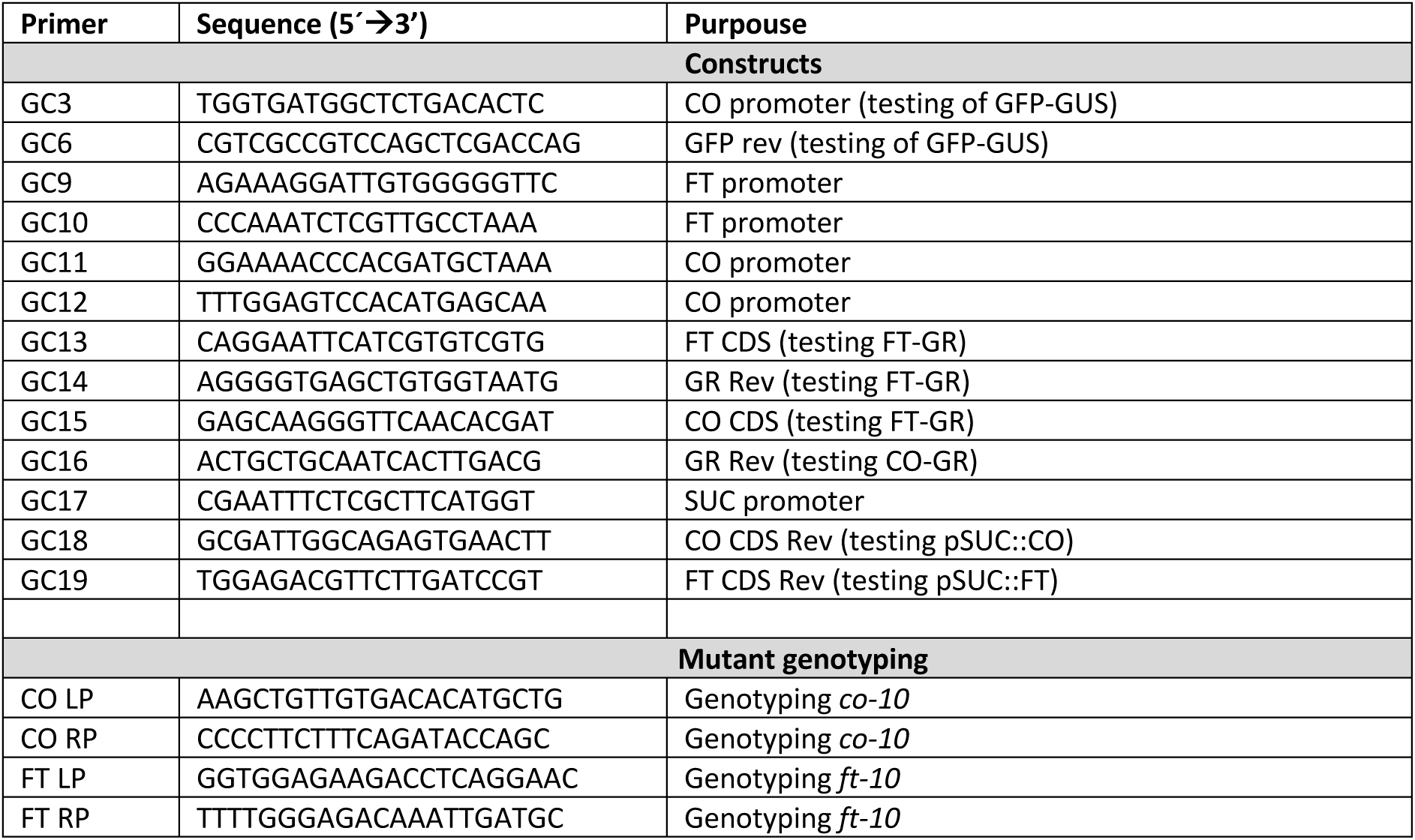
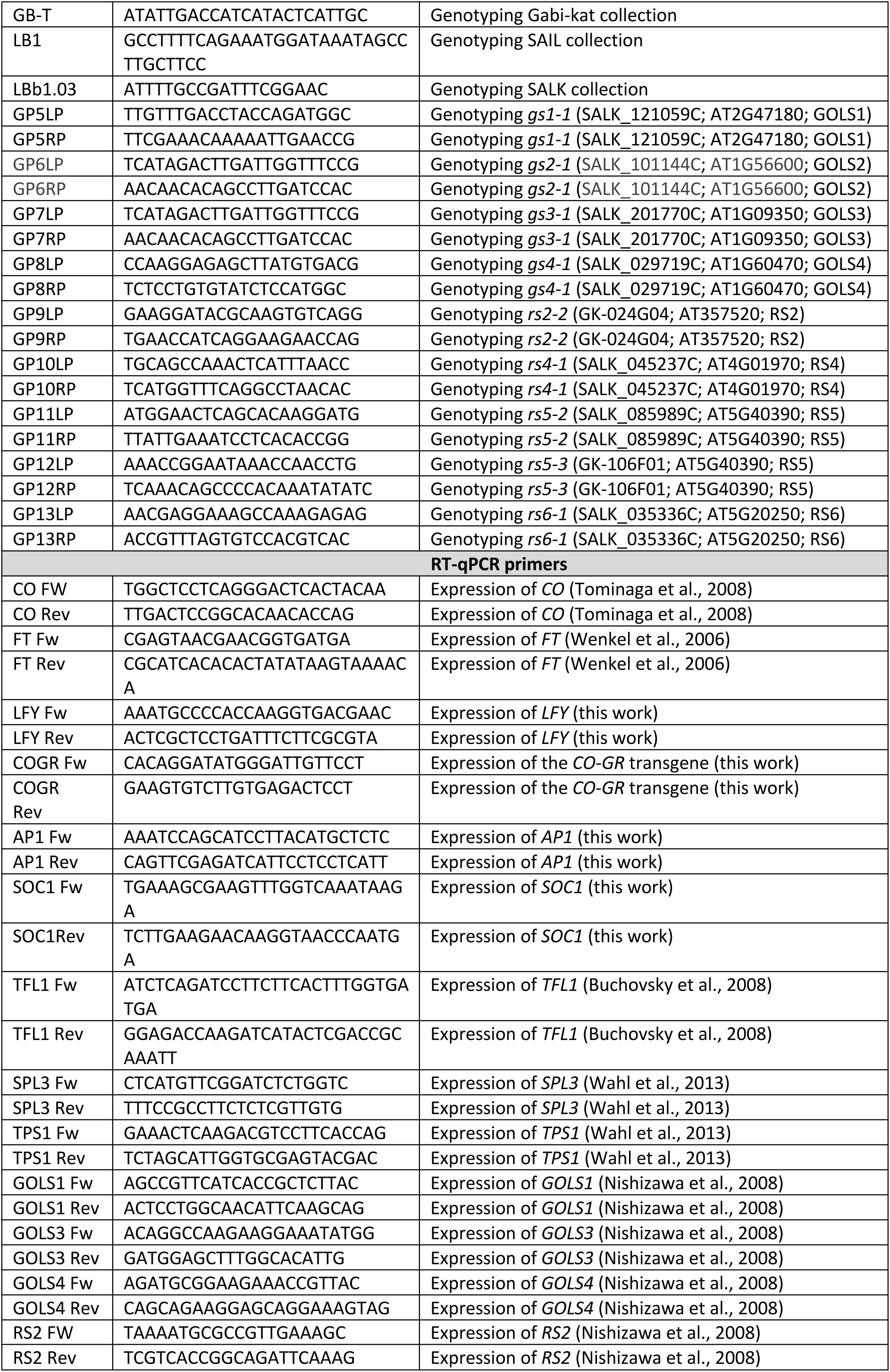
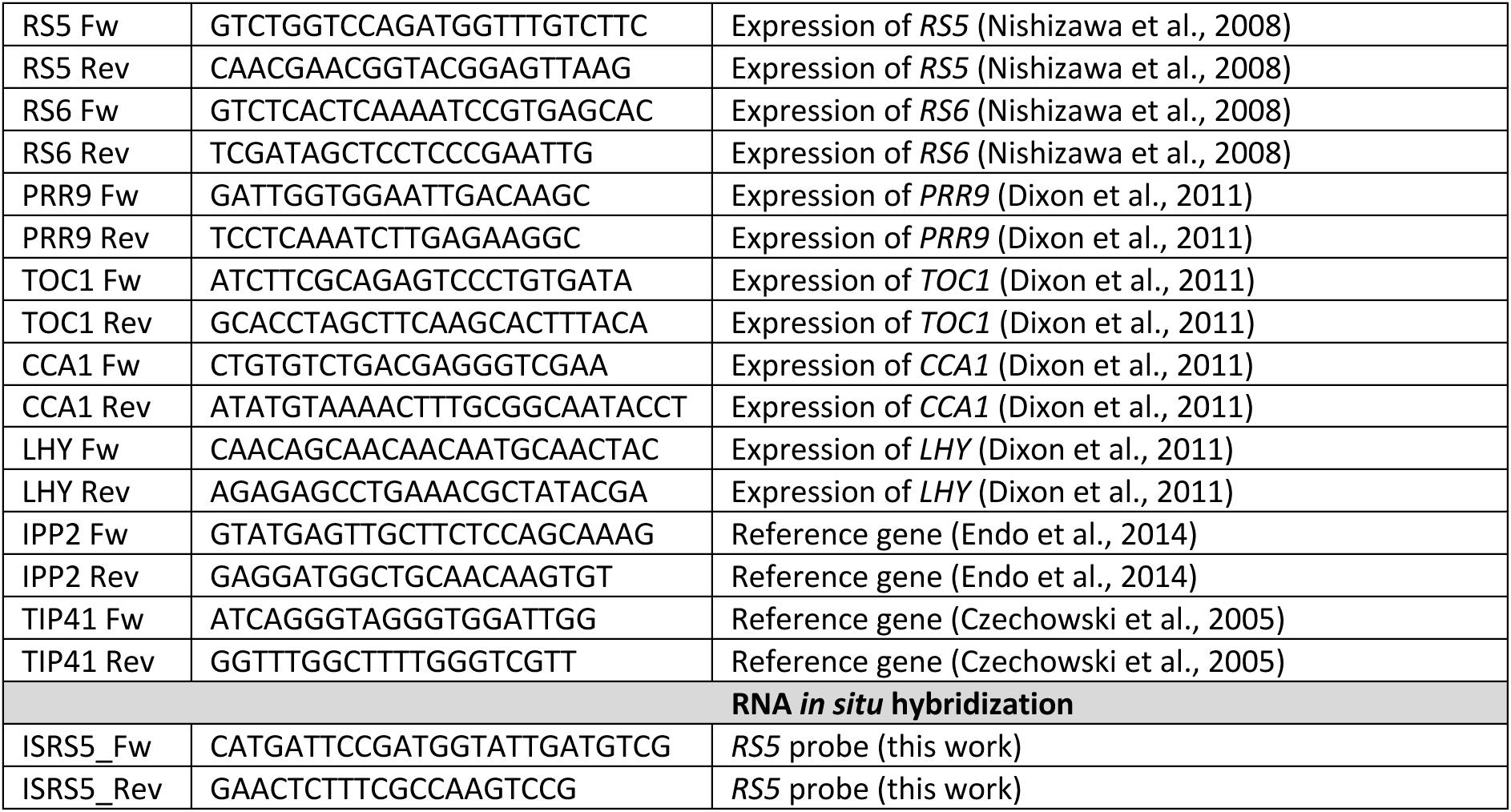
List of primers used in this work.

